# After traumatic brain injury oligodendrocytes regain a plastic phenotype and can become astrocytes

**DOI:** 10.1101/2021.06.18.448919

**Authors:** Xianshu Bai, Na Zhao, Wenhui Huang, Laura C. Caudal, Renping Zhao, Johannes Hirrlinger, Wolfgang Walz, Frank Kirchhoff, Anja Scheller

## Abstract

After acute brain injuries various response cascades are evoked that direct the formation of the glial scar. Here, we report that acute lesions associated with a disruption of the blood-brain barrier trigger a re-programming within the oligodendrocyte lineage. In PLP-DsRed1/GFAP-EGFP and PLP-EGFP_mem_/GFAP-mRFP1 transgenic mice with cortical injuries, we transiently found PLP transgene-labelled cells with activated GFAP promoter activity adjacent to the lesion site. We termed them AO cells, based on their concomitant activity of <u>a</u>stro- and <u>o</u>ligodendroglial genes. By fate mapping using PLP- and GFAP-split Cre complementation and NG2-CreER^T2^ mice we observed that major portions of AO cells surprisingly differentiated into astrocytes. Using repeated long-term *in vivo* two-photon laser-scanning microscopy (2P-LSM) we followed oligodendrocytes after injury. We observed their conversion into astrocytes via the AO cell stage with silencing of the PLP promoter and simultaneous activation of the GFAP promoter. In addition, we provide evidence that this oligodendrocyte-to-astrocyte conversion depends on local cues. At the lesion site higher expression levels of various glial differentiation factors were detected. And indeed, local injection of IL-6 promoted the formation of AO cells. In summary, our findings highlight the plastic potential of oligodendrocytes in acute brain trauma. An altered environmental milieu affects gene expression programs of mature oligodendrocytes and induces a plastic differentiation stage with astrogliogenic potential via transitional AO cells.

## Introduction

Oligodendrocytes, the myelin forming cells of the central nervous system (CNS), are terminally differentiated and originate from their precursor cells (OPCs, also termed NG2 glia) under both physiological and pathological conditions. While mammalian oligodendrocytes appear to be particularly sensitive to injuries^1^, their lower vertebrate counterparts display a more plastic behavior^2^. In the goldfish optic tract, oligodendrocytes do not only survive nerve lesion, they even dedifferentiate into elongated bipolar cells before they start to myelinate again^3^. So far, a similar dedifferentiation of rodent oligodendrocytes has been suggested by *in vitro* studies^4, 5^ and in an epigenetic analysis of MBP-Cre/loxP fate mapping^6^.

In the mammalian adult brain, the plastic behavior within the oligodendrocyte lineage is far better established for OPCs, which have consistently been found to generate astrocytes after brain injuries, while under healthy conditions only embryonic OPCs could generate astroglial cells^7, 8^. The contribution of astrocytes formed by OPCs in the injured adult brain appeared variable and strongly dependent on animal models and/or injury paradigms. OPC-derived astrocytes were detected in a variety of insults such as stab wound injury (SWI), spreading depression in grey matter (rat brain) or cryoinjury (transgenic mice)^9-12^. However, in general, varying numbers of astrocytes were found^11-13^, mostly explainable with time points of inducing the lesion pertaining to the age and time of recombination induction and varying time windows of analysis after injury.

Here, we explored the plasticity of oligodendrocytes in the mouse brain after acute cortical injuries *in vivo*. By analyzing a variety of genetically modified mouse models we provide strong evidence that not only OPCs but also mature oligodendrocytes give rise to astrocytes after acute cortical injuries via a transitional AO cell possessing astro-and oligodendroglial properties.

## Results

### Acute injury induces the generation of astrocytes from cells of the oligodendrocyte linage

To investigate the differentiation potential along the oligodendrocyte lineage, i.e. oligodendrocyte precursor cells (OPCs) and mature oligodendrocytes, to generate astrocytes *in vivo*, we employed Cre/loxP fate mapping by taking advantage of NG2-CreER^T2^ knockin mice). To label oligodendrocyte lineage cells (OLCs), we injected tamoxifen in 7-week-old NG2-CreER^T2^ x R26^fl^STOP^fl^tdTomato (R26-tdT) mice starting either 10 or 30 days before a stab wound injury (SWI, dbi), and analyzed cellular responses one week post injury (wpi) (**Fig. 1a, b**). We detected lesion-induced GFAP expression in 25.5 ± 4.4 % of tdT^+^ cells (240/931 cells, n = 3 mice) when gene recombination was induced 10 dbi (**Fig. 1c, d, f**, arrows). Over 70 % of the GFAP^+^tdT^+^ cells (71.5 ± 6.5 %, (18.4 ± 3.8 % of all tdT^+^ cells) were also PDGFRα-positive, and therefore classified as OPCs (**Fig. 1e, f**, GFAP^+^PDGFRα^+^tdT^+^, triangles, 175/931 cells, n = 3 mice). We regarded the remaining 28.5 ± 6.5 % % of recombined and PDGFRα-negative cells as *bona fide* astrocytes (**Fig. 1e, f**, GFAP^+^PDGFRα^-^ tdT^+^, arrowheads, (7.1 ± 1.0 % of total tdT^+^ cells, 65/931 cells). Their identity could be further substantiated by glutamine synthetase (GS) immunoreactivity and by a typical astrocytic morphology with fine, highly arborized processes and contacts to blood vessels (**Fig. 1g**). The percentage of astrocytes among all recombined tdT^+^ cells was strongly increased when analyzed four weeks after injury (7.1 ± 1.0 % to 19.6 ± 0.8 % (78/397 cells)). Under physiological, non-injury control conditions, no astrocytes were generated from OPCs, as described before^7, 8, 14^.

**Figure 1.**
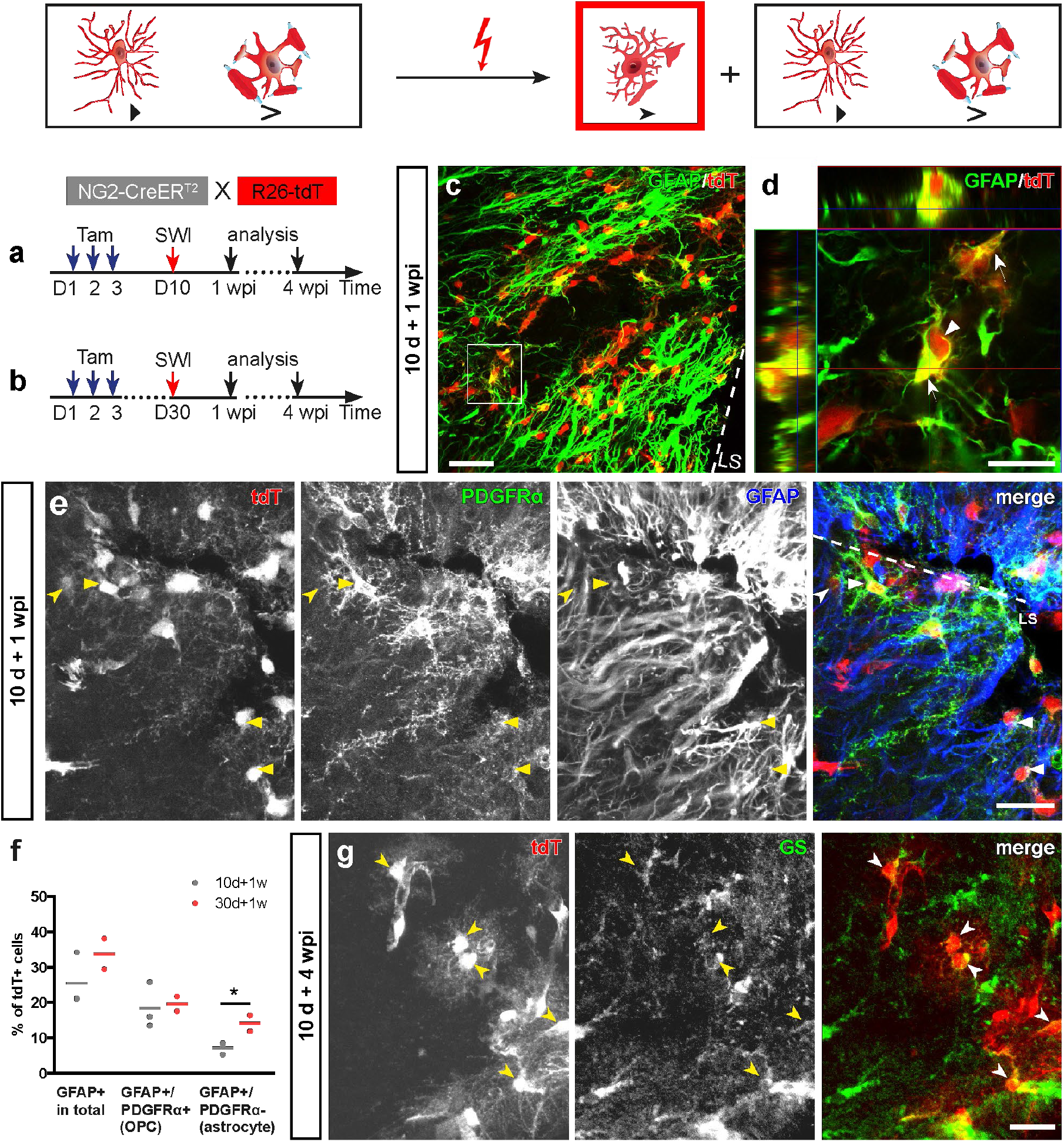
Oligodendrocyte lineage cells give rise to astrocytes after stab wound injury (SWI). **a, b**, Scheme of DNA recombination induction and analysis of NG2-CreER^T2^ mice. **c**, Intensive GFAP expression in recombined cells in NG2-CreER^T2^ mice 1 wpi. **d**, Orthogonal projection of a GFAP expressing recombined cell. **e**, GFAP expression of OPCs (PDGFRα^+^tdT^+^, triangles) and astrocytes (PDGFRα^-^tdT^+^, arrowheads) adjacent to the injury 1 wpi. **f**, Quantification of recombined cells 1 wpi showed an increased number of GFAP^+^ recombined astrocytes after a 30-day period left for gene recombination, compared to a 10-day period. **g**, Four weeks after SWI, GS^+^tdT^+^ astrocytes (arrowheads) could still be detected adjacent to the lesion. ^*^: p < 0.05, two tailed unpaired t-test. LS: lesion site, triangles: OPCs, arrowheads: astrocytes. Scale bars in **c** = 50 µm, **d** = 10 µm, and **e, g** = 25 µm.

When leaving 30 d instead of 10 d between tamoxifen injection and SWI (**Fig 1b**), the proportion of astrocytes (GFAP^+^tdT^+^PDGFRα^-^) was further increased and twice as large (**Fig. 1f**, 14.1 ± 2.3 % (76/527 cells, n = 3 mice) vs. 7.1 ±1.0 %, p=0.0445), while that of OPCs remained constant (**Fig. 1f**, GFAP^+^tdT^+^PDGFRα^+^, 19.6 ± 2.1 % (105/527 cells) vs. 18.4 ± 3.8 %). The longer time period left for gene recombination (30 dbi compared to 10 dbi) the more time for lineage progression, and thereby increased the relative quantity of labeled oligodendrocytes (20.2 % vs 33.9 %)^8^, in line with other studies^14, 15^. Hence, the higher number of newly generated astrocytes was correlated with the higher percentage of recombined tdT^+^ oligodendrocytes. Thereby, these data provided the first hint that not only OPCs, but also a subpopulation of mature oligodendrocytes might generate astrocytes after SWI.

To further substantiate and directly visualize the activation of the GFAP gene in different glial cell stages, we bred the NG2-CreER^T2^ x R26-tdT mice to a GFAP-EGFP_GFEC_mouse line with expression of the green fluorescent protein in astrocytes (**Supplementary Fig. 2a-e**, asterisks). And indeed, beside numerous OPCs (tdT^+^EGFP^+^PDGFRα^+^, 51.6 ± 2.5 %, 97/194 cells, n=3 mice), we also observed PDGFRα-negative double-fluorescent cells (tdT^+^EGFP^+^PDGFRα^-^, 48.4 ± 3.0 %, 97/194 cells) at 1 wpi with a roundish cell body and few fine processes. After four weeks, we could identify double-fluorescent reporter^+^ cells, now immune-positive for GFAP (4 wpi, **Supplementary Fig. 2f-h**).

Although these data, based on inducible Cre/loxP recombination and cell-specific EGFP expression, already suggested that not only OPCs, but also oligodendrocytes could generate astrocytes, these experiments did not allow an unequivocal discrimination of astrocytes generated from OPCs or derived from oligodendrocytes.

### Astroglial differentiation from oligodendrocytes after acute trauma

To further test the origin of astrocytes directly from oligodendrocytes, we used transgenic split-Cre mice that permanently label the newly formed cells by Cre complementation in GFAP-N-terminal (NCre) and PLP-C-terminal (CCre) split-Cre transgenic mice. Using immunohistochemical analysis of healthy adult GFAP-NCre_GCNT_x PLP-CCre_PCCK_mice selected from various transgenic founders, CCre expression could only be detected in mature oligodendrocytes (**Fig. 2a**, >95 % of GSTπ^+^, 116/122 cells; **Fig. 2b**, MOG), but not in OPCs (**Fig. 2c**; NG2^+^; **Fig. 2d**, PDGFRα^+^). After cortical injury we found strong reporter expression indicating efficient split Cre complementation (**Fig. 2f, Supplementary Fig. 3a-c, e, f**), which was absent in the uninjured cortex (**Fig. 2e** and **Supplementary Fig. 3c, d**). Simultaneously, we did not only detect GFAP expression, but we also observed a typical astroglial morphology in these cells derived from PLP-CCre expressing oligodendrocytes (**Fig. 2g**, GSTπ^+^), (**Fig. 2h**, GFAP^+^tdT^+^CCre). The generation of GFAP^+^tdT^+^ cells could also be induced in PLP-CreER^T2^ transgenic mice, but we were not able to detect other cells than astrocytes in GFAP-CreER^T2^ mice (**Supplementary Fig. 3g-i**). These data provide strong genetic evidence that astrocytes can be generated from oligodendrocytes after SWI.

**Figure 2.**
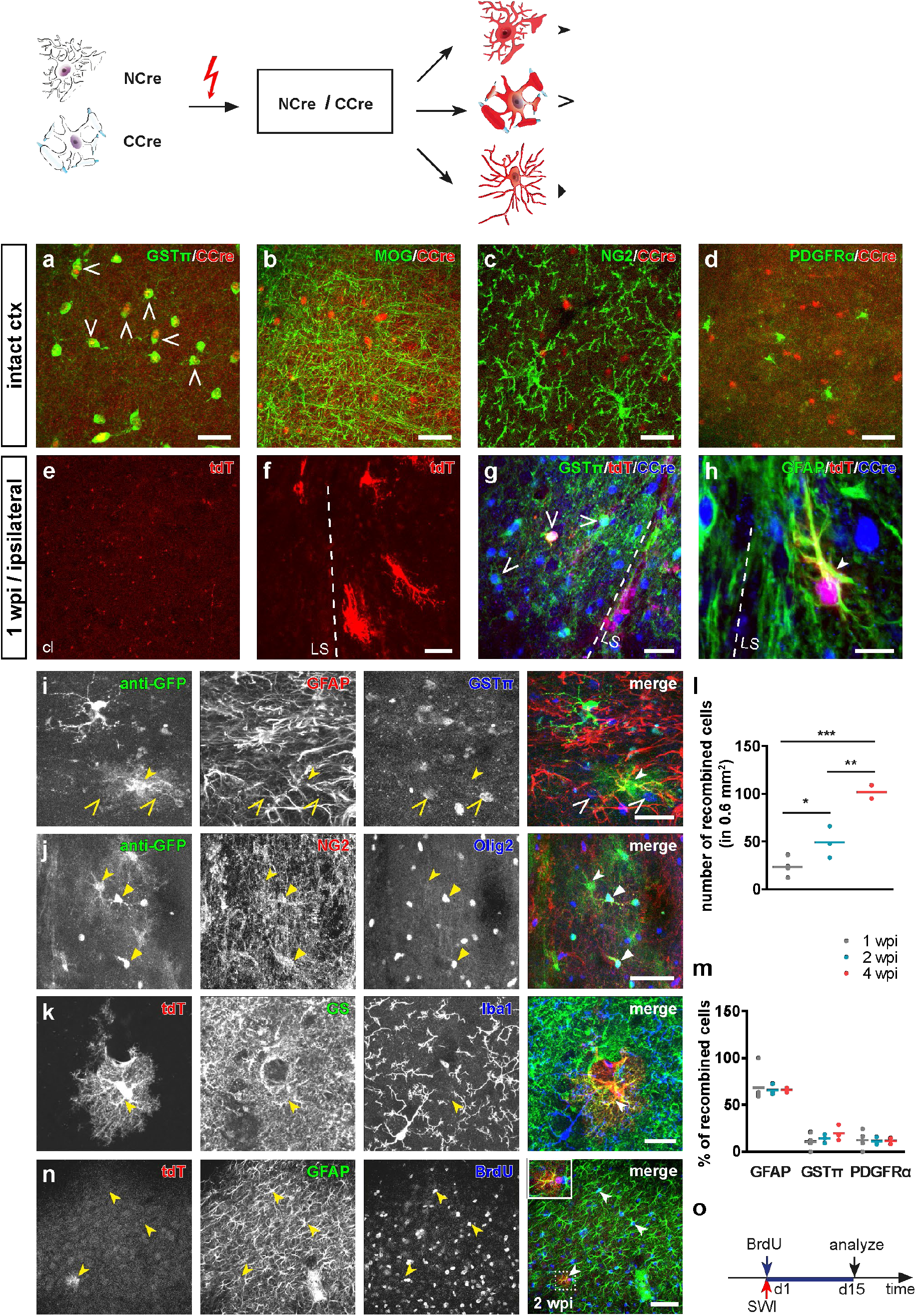
Split-Cre complementation uncovers the potential of mature oligodendrocytes to generate astrocytes after SWI. **a-d**, The split-Cre C-terminal fragment (CCre) was exclusively detectable in mature oligodendrocytes in the intact cortex (GSTπ^+^, **a**; MOG^+^, **b**), but not in NG2 glia (NG2^+^, **c**; PDGFRα^+^, **d**). **e, f**, No recombination was observed in the intact cortex (**e**) but adjacent to the lesion site (LS, **f**). **g, h**, CCre expression in recombined oligodendrocytes (GSTπ^+^tdT^+^, **g**) and astrocytes (GFAP^+^tdT^+^, **h**) at LS 1 wpi. **i-k**, Immunolabeling revealed that recombined cells could give rise to astrocytes (GFAP^+^, **i**; GS^+^, **k**), oligodendrocytes (GSTπ^+^, **i**) and OPCs (NG2^+^, **j**), but not to microglia (Iba1^+^, **k**). **l, m**, The number of total recombined cells increased with time after SWI (1, 2 and 4 wpi) at the lesion site (0.6 mm^-2^), while the proportion of recombined cell types did not change over time (**m**), being mainly astrocytes. **n**, BrdU was incorporated by recombined astrocytes, indicating proliferative capacity of recombined astrocytes. **o**, Experimental schedule for BrdU administration. ^*^: p<0.05, ^***^: p<0.001, compared with 1 wpi; ^**^: p<0.01, compared with 2 wpi, one-way ANOVA. Scale bars in **a-g, i, j** = 25 µm, **h, k** = 10 µm, **n** = 50 µm.

Quantification of recombined cells in split-Cre mice at the lesion site (about 0.6 mm^2^ area of the 40-µm, frontal brain sections) (**Fig. 2i-k**) showed an increase in total number of recombined cells over time (**Fig. 2l**, 1 wpi (23.5 ± 4.9 cells/0.6 mm^2^), 2 wpi (49.0 ± 9.5 cells/0.6 mm^2^, p = 0.049) and 4 wpi (102 ± 7.0 cells/0.6 mm^2^, p = 0.0008 vs. 1 wpi and p = 0.028 vs. 2 wpi), n = 3 mice,). However, the ratios between the glial cell types did not change during the time period of analysis (**Fig. 2m**). The presumed astrocytes represented the majority of recombined cells (**Fig. 2i, k, m**, 2 wpi: 66.7 ± 0.7 % (190/294 cells, n = 3), GFP^+^GFAP^+^GSTπ^-^, tdT^+^GS^+^Iba1^-^, tdT^+^GFAP^+^So×10^−^). The remaining cells are largely oligodendrocytes (**Fig. 2i, m**, 2 wpi: 14.7 ± 2.7 % (49/336 cells, n = 3), tdT^+^GSTπ^+^) or OPCs (**Fig. 2j, m**, 2 wpi: 11.6 ± 0.4 % (30/261 cells, n = 3), tdT^+^PDGFRα^+^ or GFP^+^NG2^+^) at equal amounts. These split-Cre data suggest that oligodendrocyte-derived recombined cells become astrocytes within four weeks.

To investigate whether also oligodendrocyte-derived astrocytes respond to injury with proliferation, we performed a BrdU assay in split-Cre mice (**Fig. 2n, o**). Among the BrdU^+^tdT^+^ cells (71.7 ± 7.8 %, 300/415 cells, n = 3), the majority were astrocytes (**Fig. 2n**, GFAP^+^, 80.6 ± 2.3 %, 114/141 cells, n = 3). Therefore, similar to *bona fide* reactive astrocytes, also oligodendrocyte-derived astrocytes proliferate in response to injuries.

In split-Cre mice we detected recombined cells distributed in a cortical layer-dependent gradient with the highest number of newly generated astrocytes in the upper cortical layers, and less next to the corpus callosum (**Supplementary Fig. 4a-e and Supplementary table 1**). In contrast, newly generated oligodendrocytes were mainly observed next to the corpus callosum but less under the pia, displaying a similar distribution pattern like recombined oligodendrocytes in non-lesioned NG2-CreER^T2^ mice. These results indicate that the intrinsic properties of local niches appear unaffected of lesion size or transgenic mouse model and that oligodendroglial differentiation is preferred in the area near corpus callosum. (**Supplementary Fig. 4f-j** and **Supplementary table 1**).

### Injured oligodendrocytes generate a plastic cell type distinct from OPCs

Since the split-Cre fate mapping provided genetic evidence only for the endpoint of astrocyte generation from injured oligodendrocytes, we employed PLP-DsRed1/GFAP-EGFP_GFEA_double fluorescent mice to directly observe and characterize cells at the transitional stage towards the generation of astrocytes (**Fig. 3a**). The transition was observed based on the simultaneous expression of DsRed1 and EGFP, indicating the coincident activity of PLP and GFAP promoters. These transitional cells (with overlapping <u>a</u>stro- and <u>o</u>ligodendroglial properties) were termed AO cells. And indeed, directly adjacent to the lesion site intensive astro- and oligodendroglial promoter activation could be observed (**Fig. 3a, c**, at 13 dpi) and numerous AO cells could be identified already 3 dpi (**Fig. 3c**, asterisks). Such fluorescently labelled cells, however, were never detected in the intact cortex (**Fig. 3b**). AO cells displayed round somata (**Fig. 3d**, diameter around 10 µm) with few processes (**Fig. 3d**, arrows) and DsRed1 protein aggregates at all observed time points between 3 dpi and 14 dpi (**Supplementary Fig. 5a)**: Such aggregates are a characteristic property of many reef coral fluorescent proteins such as DsRed1 indicating a long-term expression^16^. Here, these DsRed1 aggregates imply a start of expression prior to the injury, i.e. in uninjured PLP-DsRed1-positive oligodendrocytes. In a volume of 2.4 × 10^−2^ mm^3^ at the lesion site, about 50 AO cells were found (21 cells/1×10^−2^mm^3^). Please note, only a minority (23.5 %) of all putative AO cells with coincident promoter activity could be recognized by simultaneous expression of EGFP and DsRed1 due to the limited penetrance of the transgenes (only 42.7 % of astrocytes express the GFAP-EGFP and 55.1 % of oligodendrocytes express the PLP-DsRed1 transgenes, **Supplementary Fig. 6a-f**), while only few OPCs express DsRed1 either ipsi-or contralateral (2.1 % and 1.7 % respectively, **Supplementary Fig. 6g-i**). As expected AO cells were positive for oligodendrocyte lineage markers (**Fig. 3e**, Sox10, 96 ± 3 %, 133/138 cells, n = 3 mice; **Supplementary Fig. 7a**, Olig2, 93 ± 2 %, 83/89 cells, n = 3 mice). We could also observe quite a few AO cells expressing the mature oligodendrocyte marker GSTπ (**Supplementary Fig. 7b**, 2 %, 3/143 cells). However, AO cells were not expressing other astroglial (**Fig. 3f**, Camsap1; **Supplementary Fig. 7d**, GFAP), stem cell (**Fig. 3g**, Sox2), OPC (**Supplementary Fig. 7e**, PDGFRα), neuronal (**Supplementary Fig. 7f**, NeuN) or microglial markers (**Supplementary Fig. 7g**, Iba1). These data indicate that AO cells belong to the oligodendrocyte lineage but are different from OPCs as well as from stem cells or astrocytes.

**Figure 3.**
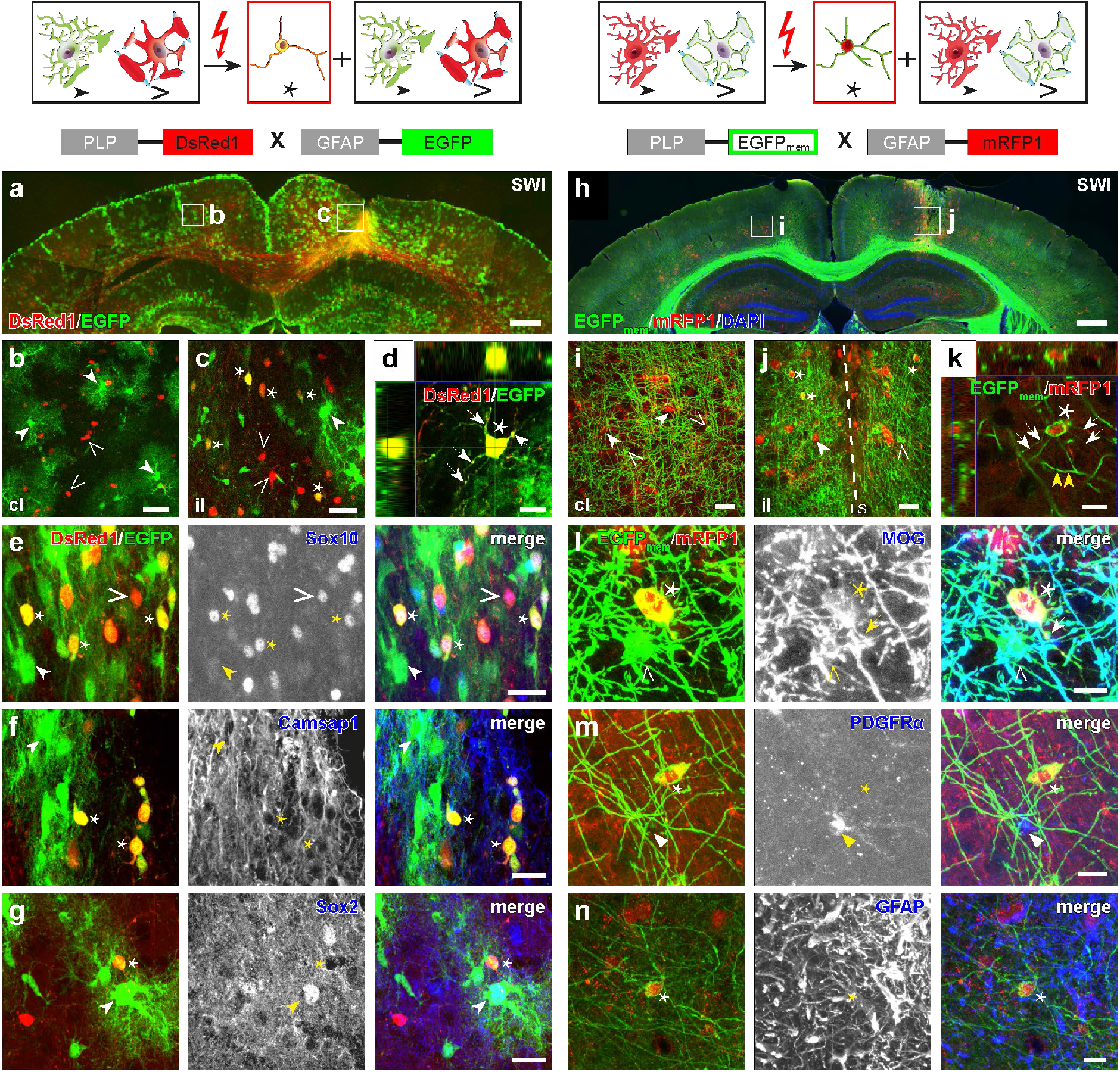
Acute injuries induce a novel transient stage of oligodendrocyte lineage cells, AO cells. **a, h**, Intensive glial reaction close to the lesion site (SWI) in cortex at 13 dpi of PLP-DsRed1_PRDB_/GFAP-EGFP_GFEA_ (**a**) and PLP-EGFP_mem_/GFAP-mRFP1 (**h**) mice. **b, c**, Magnified views of regions indicated in **a. i, j**, Magnified views of regions indicated in **h**. Numerous AO cells expressed either EGFP^+^DsRed1^+^ (asterisks in **c**) or EGFP _mem_^+^ mRFP1^+^ (asterisks in **j**) and were located close to the lesion site, but not at the contralateral side (**b and i**). **d, k**, The orthogonal projection demonstrated unequivocal co-expression of EGFP/DsRed1 (**d**) or EGFP_mem_/mRFP1 (**k**) in AO cell somata (asterisk) and thin processes (white arrows). Yellow arrows indicate myelin of adjacent oligodendrocytes. **e-g**, Immunostaining revealed that EGFP^+^/DsRed1^+^ AO cells expressed the oligodendrocyte lineage marker Sox10 (**e**), but neither the astrocyte lineage marker Camsap1 (**f**), nor the stem cell marker Sox2 (**g**). **l-n**, Immunostaining revealed that EGFP _mem_^+^ mRFP1^+^ AO cells expressed the mature oligodendrocyte marker MOG (**l**), but neither the OPC marker PDGFRα (**m**), nor the astrocyte marker GFAP (**n**). Open triangles: oligodendrocytes, arrowheads: astrocytes, asterisks: AO cells. Scale bars in **a, h** = 500 µm; **b, c, i, j** = 25 µm, **d, k-n** = 10 µm, **e-g** =20µm.

To further confirm oligodendroglial properties of AO cells, we used a complementary approach by analyzing the double transgenic mouse line, PLP-EGFP_mem_/GFAP-mRFP1 (**Fig. 3h**), which uses exactly the same promoters (PLP/GFAP) to drive expression. But, here membrane-bound EGFP labels oligodendrocytes, while the red mRFP1 is expressed in astrocytes under physiological conditions (**Fig. 3i**). In these PLP-EGFP_mem_ mice all EGFP-expressing cells are mature oligodendrocytes (**Supplementary Fig. 8a, b**, 100 %, GSTπ^+^). From 2 dpi on, similar to PLP-DsRed1/GFAP-EGFP mice, we observed double-labelled AO cells expressing membrane-bound EGFP and cytosolic mRFP1 (**Fig. 3j, k**, asterisk/arrows). Similar to DsRed1 also its mutant form mRFP1 frequently forms aggregates after longer expression periods. This was very apparent in the adjacent astrocytes (**Supplementary Fig. 5b**, arrows). In contrast, a uniform distribution of cytosolic mRFP1 was found in AO cells, thereby indicating a short time of GFAP promoter activity (**Supplementary Fig. 5b**, asterisk). In PLP-EGFP_mem_/GFAP-mRFP1 mice, we could detect several marker proteins for myelin on AO cells with EGFP expression in the membrane (**Fig. 3l**, MOG; **Supplementary Fig. 8c, d**, MAG, PLP, respectively), but never the OPC marker PDGFRα (**Fig. 3m**). Despite GFAP promoter activity in AO cells, the protein itself could never be detected (**Fig. 3n**). Since the GFAP gene is very sensitive to pathological alterations, we asked whether a completely different astroglial gene, the glutamate/aspartate transporter GLAST, would also be activated in AO cells. For that purpose, we crossbred PLP-EGFP_mem_, mice with the astrocyte-specific knockin mouse GLAST-CreER^T2^ x R26-tdT. Indeed, we observed tdT^+^ and EGFP^+^ cells at the lesion site. Since these cells were also immune-positive for GFAP, we regarded them as astrocytes generated from AO cells as well (**Supplementary Fig. 8e-g**).

These results further confirm our notion that AO cells originate from oligodendrocytes, are different from OPCs, can activate astrocyte-specific genes and can change their fate to the astroglial lineage.

### Whole-cell membrane currents of AO cells are highly variable

To characterize physiological properties of AO cells, in addition to marker protein expression, we performed whole-cell patch-clamp recordings and tested for the patterns of membrane currents that are characteristic for OPCs, oligodendrocytes and astrocytes. After SWI of PLP-DsRed1/GFAP-EGFP mice at P20 (**Fig. 4a**, 3 dpi to 4 dpi), AO cells (**Fig. 4b**) were recorded with a KCl-based intracellular solution and held at −80 mV. Whole-cell membrane currents of AO cells were dominated by K^+^ currents, lack of voltage-gated Na^+^ currents and were, thereby, very typical for glia. However, individual AO cells displayed a high variability in respect to the presence of voltage-gated, outwardly rectifying K^+^ currents, symmetrical non-rectifying or inwardly rectifying K^+^ currents (**Fig. 4c-f**). Furthermore, we compared whole-cell currents and their membrane properties in identified OPCs (NG2-EYFP), oligodendrocytes (PLP-DsRed1) and astrocytes (GFAP-EGFP) under physiological, non-injury conditions (without SWI, cl) and in their activated state after SWI (il). AO cells typically exhibited a slightly more positive resting membrane potential (V_m_=-67.8 ± 4.2 mV, n = 18 cells) than OPCs (**Fig. 4g**, cl: V_m_=-79.5 ± 1.4 mV, n = 14 cells, p=0.0235; il: V_m_= −81.1 ± 1.6 mV, n = 11 cells, p=0.0232), but not from those of astrocytes or oligodendrocytes. AO cells had a higher membrane resistance (R_m_=92 ± 10 MΩ) than astrocytes (**Fig. 4h**, R_m_=33 ± 2 MΩ, n = 21 cells (cl); R_m_=68 ± 9 MΩ, n = 12 cells (il), p=1.67E-06), but not in comparison to oligodendrocytes (R_m_=60 ± 5 MΩ, n = 11 cells (cl); R_m_=95 ± 8 MΩ, n = 13 cells (il)) or OPCs (R_m_=63 ± 6 MΩ (cl); R_m_=91 ± 10 MΩ (il)).

**Figure 4.**
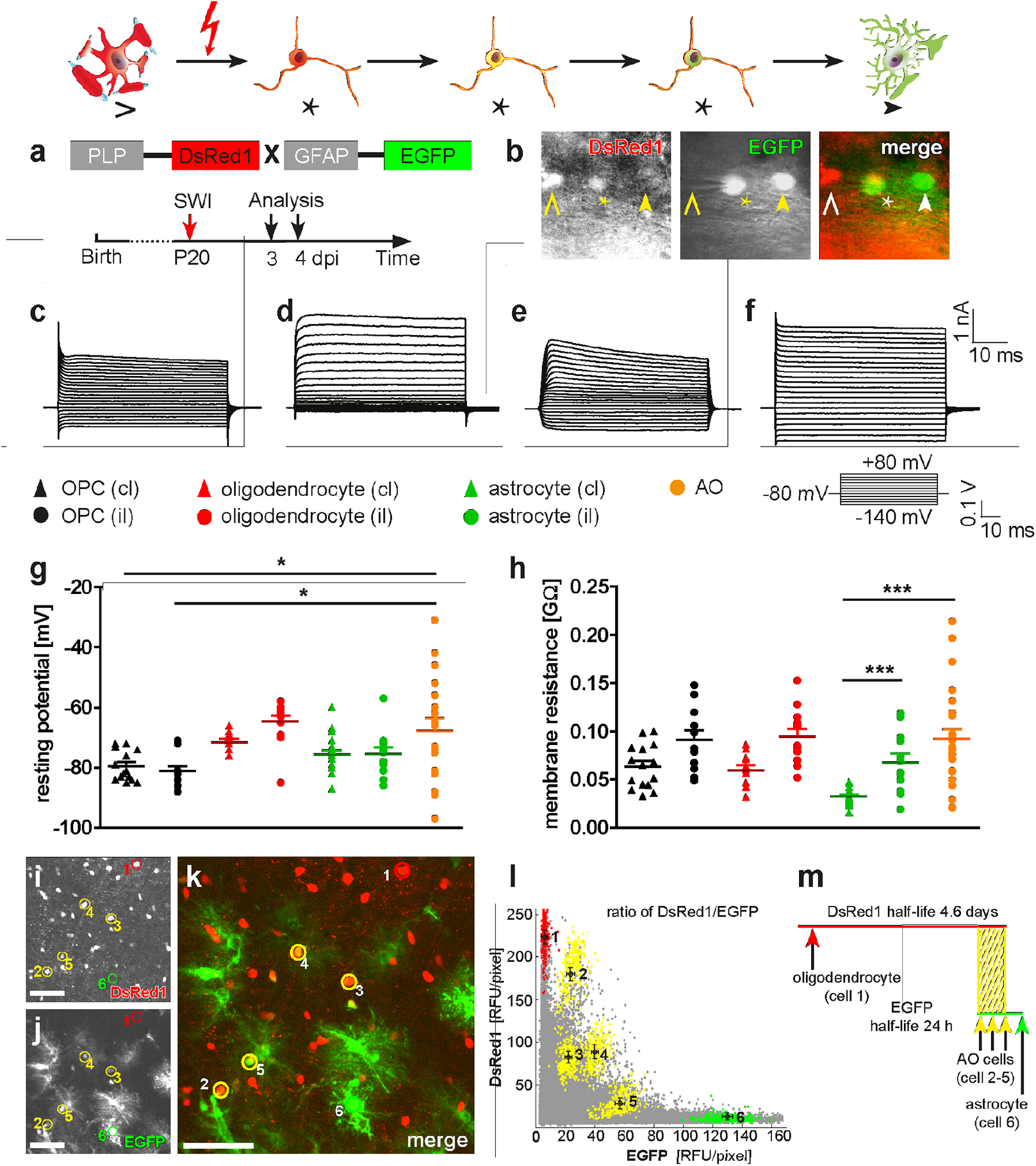
AO cells display variable electrophysiological properties and transgene expression. **a**, Time schedule of experiment. **b**, AO cell (EGFP^+^DsRed1^+^, asterisk) identified next to an astrocyte (EGFP^+^DsRed1^-^, arrowhead) and oligodendrocyte (EGFP^-^DsRed1^+^, open triangle). **c-f**, AO cells with different membrane properties. **g**, Comparison of resting membrane potentials showed larger variability among AO cells compared to other glial cell types in contra-and ipsilateral. **h**, Analysis of membrane resistances showed broad variability in AO cells with significant differences to astrocytes of the contralateral side, but not of AO cells and activated glial cells at the injury side. **i-l**, Quantification of different expression ratios of DsRed1/EGFP in AO cells ranging from orange (DsRed1 > EGFP) and yellow (DsRed1 = EGFP) to green (DsRed1 < EGFP) (cell *2-5* in **k**) fluorescence. Oligodendrocytes (cell *1* in **j**) and astrocytes (cell *6* in **i**) serve as controls. **m**, Temporal overlap of fluorescent protein expression in glial cells as suggested by their half-life. PLP-promoter controlled DsRed1 (with half-life of ∼4.6 days) is still present when acute injuries activate EGFP expression in AO cells (half-life ∼24 h)^17^. Open triangle: oligodendrocyte, arrowhead: astrocyte, asterisk: AO cell. ^*^: p < 0.05, ^**^: p < 0.01, ^***^: p < 0.001, compared with AO cells, one-way ANOVA. Scale bars in **i-k** = 50 µm.

The high variability of electrophysiological properties among individual AO cells does not allow them to be classified as oligodendrocytes, astrocytes or OPCs, but rather suggests a unique glial cell type with transitional properties. Indeed, a similar variability we observed at the level of transgene expression. AO cells in the lesioned area displayed a broad and variable range of EGFP and DsRed1 levels (i.e EGFP/DsRed1 ratio). Cells with high DsRed1 expression were still assigned to the oligodendrocyte lineage, while higher EGFP expression indicated a more astroglial phenotype (**Fig. 4i-l**). When AO cells were formed from oligodendrocytes, the transgenic GFAP promoter would be activated and the PLP promoter activity (the endogenous as well as the transgenic) would decrease and, subsequently, transcription would stop. Since the half-life of DsRed1 is 4.6 days^17^, DsRed1 was still detectable in AO cells when the PLP promoter activity had ceased (**Fig. 4m**).

These results indicate that AO cells are cells in transition, thereby explaining their variability of membrane properties and transgene expression levels.

### *In vivo* two-photon laser scanning microscopy (2P-LSM) visualizes the conversion of oligodendrocytes to astrocytes directly

To directly investigate the change from oligodendrocyte to astrocyte, we performed *in vivo* 2P-LSM. In PLP-DsRed1/GFAP-EGFP mice, we could detect AO cells already at 3 dpi (**Fig. 5a-c**, cells *2, 3*) remaining in this stage for the next days. We show exemplarily the tracing of a single oligodendrocyte (**Fig. 5a**, DsRed1^+^EGFP^-^, cell *1*, open triangle) with starting EGFP-expression 3 dpi and turning into an AO cell (DsRed1^+^EGFP^+^) 5 dpi (**Fig. 5b**, cell *1*). Again DsRed1 aggregates could be found in processes (**Fig. 5b**, arrows), confirming a prolonged DsRed1 expression^16^. At 6 dpi, this AO cell had stopped to express DsRed1 with subsequent protein degradation, but continued EGFP expression (**Fig. 5c**, cell *1*, arrowhead). In the same region we found other AO cells with increased levels of EGFP (**Fig. 5b**, 5 dpi, cell *2, 3*) and low-levels of DsRed1, recognizable by a higher EGFP fluorescence compared to DsRed1 at 6 dpi (**Fig. 5c, Supplementary movie 1**). Repeated observation of AO cells showed their potential to generate astrocytes. However, we could also observe AO cells (one example given in **Supplementary Fig. 9a, b**, 6 dpi, asterisk) switching off the GFAP-EGFP transgene and turning back into a DsRed1^+^ cell, i.e. an oligodendrocyte, observed after 4 wpi (**Supplementary Fig. 9c, d**, open triangle). We did never observe an EGFP expressing astrocyte that activated the PLP-DsRed1 transgene.

**Figure 5.**
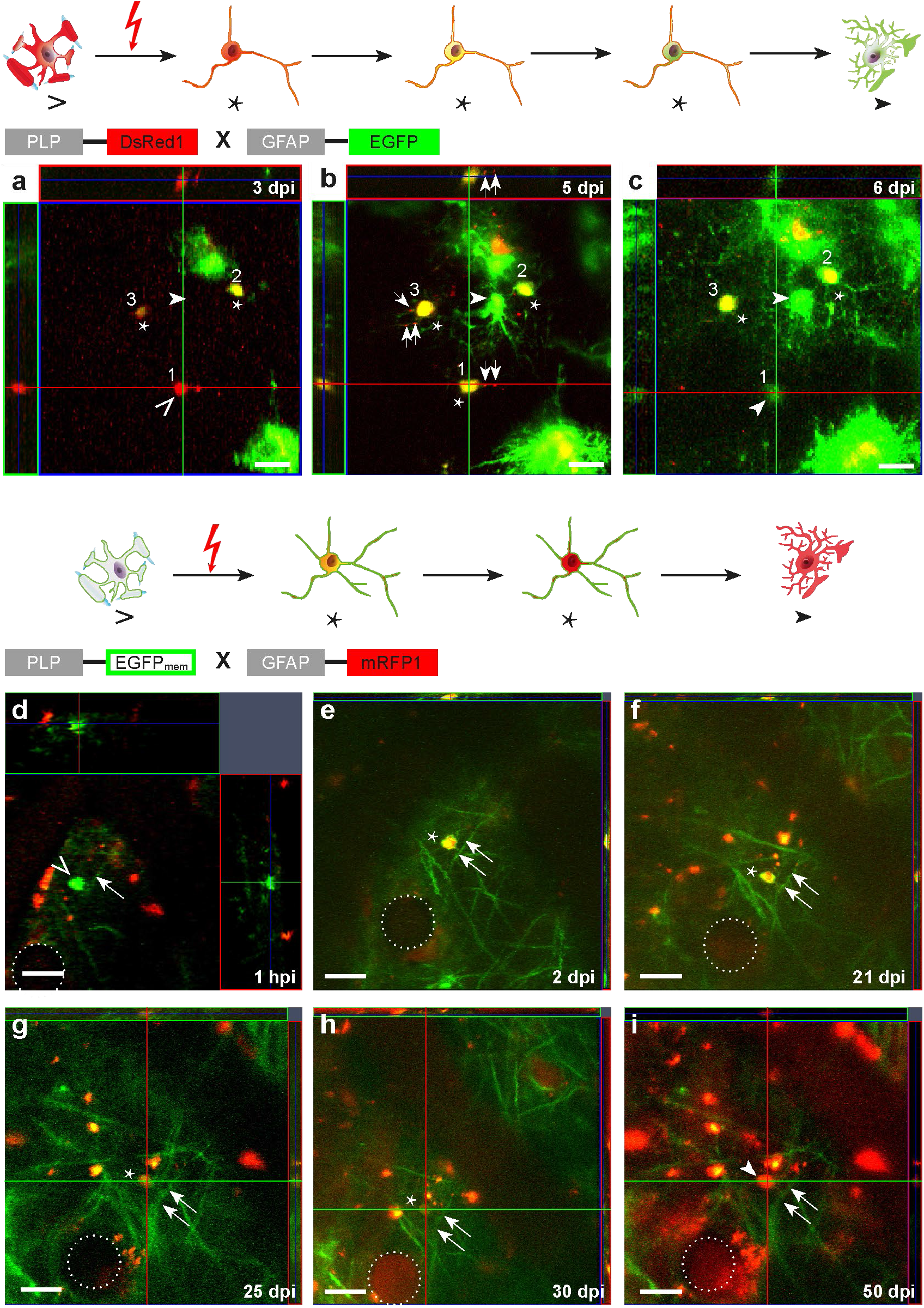
*In vivo* 2P-LSM visualizes formation of AO cells from mature oligodendrocytes and their differentiation fate. **a-c**, *In vivo* imaging of AO cells in PLP-DsRed1/GFAP-EGFP mice. **a**, Oligodendrocyte (*1*, open triangle) and AO cells (*2, 3*, asterisks) detected three days after SWI. **b**, An oligodendrocyte turned into an AO cell (*1*, asterisk) while two AO cells (*2, 3*, asterisk) stayed at the AO cell stage, but varied their DsRed1/EGFP ratios. Note the appearance of an EGFP-expressing astrocyte (arrowhead in **b**). **c**, AO cells (*1, 2, 3*) down-regulated DsRed1 expression and a single cell expressed EGFP only (*1*, arrowhead) 6 dpi. **d-i**, *In vivo* imaging of AO cells in PLP-EGFP_mem_/GFAP-mRFP1 mice. An oligodendrocyte (open triangle) was detected 1 hpi (**d**) that started to express mRFP1 (asterisks) from 2 dpi till 30 dpi (**e-g**), but with variable EGFP/mRFP1 ratios. **i**, AO cell became an astrocyte (mRFP1+/EGFP-) 50 dpi. Note that membrane-bound EGFP_mem_ (arrows in **d-i**) was still detectable 50 dpi in a newly differentiated astrocyte characterized by long-term GFAP promoter activity. Open triangle: oligodendrocyte, arrowhead: astrocyte, asterisk: AO cell. Scale bars = 20 µm.

These observations could be confirmed in a second transgenic mouse line, in PLP-EGFP_mem_/GFAP-mRFP1 mice. An oligodendrocyte (**Fig. 5d**, EGFP^+^mRFP1^-^, open triangle) started to express mRFP1 2 dpi (**Fig. 5e**, asterisk) and expressed both fluorescent proteins still 30 dpi (**Fig. 5f-h**, asterisks). But only mRFP1 expression remained 50 dpi (**Fig. 5i**, arrow head; **Supplementary movie. 2**). We also observed oligodendrocytes becoming AO cells and going back to EGFP _mem_^+^ oligodendrocytes (see example in **Supplementary Fig. 9e-g**). These results were affected by the longer half-life of membrane-bound EGFP in comparison to the cytosolic EGFP, like other membrane-confined proteins^18^. We never found mRFP^+^ astrocytes that activated the PLP-EGFP_mem_ gene. In addition, we never observed hints for phagocytosis in astrocytes, neither in oligodendrocytes or AO cells of both mouse lines. In total, we could follow 40 oligodendrocytes (of 122 in n = 17 mice) to convert via the AO cells into either astrocytes or become oligodendrocytes again. In contrast to split-Cre mice with 67 % of AO cells becoming astrocytes, by *in vivo* 2P-LSM we observed only 5 % (2 cells) becoming transgene-expressing astrocytes. Unfortunately, several AO cells could not be followed over a longer time period due to a loss of cranial window clarity, impaired light transmission and increased light scattering of the developing glial scar.

The long-term, repeated *in vivo* imaging data from two distinct transgenic mouse lines confirm that transitional AO cells originate from oligodendrocytes and either stay in the oligodendrocyte lineage or convert to astrocytes.

### Generation and differentiation of AO cells is influenced by cytokines

To test how common the formation of AO cells is, we performed two other injury models: pial vessel disruption (PVD, representing a small hemorrhagic, arterial vessel stroke) and transient middle cerebral artery occlusion (MCAO, representing a reversible large vessel stroke, **Fig. 6a**). We observed AO cells adjacent to lesion sites of both insults (**Fig. 6b**, 1 w after PVD; **Fig. 6c**, 3 d after MCAO). Therefore, we concluded that acute cortical injuries in general induce the formation of AO cells. A common feature of the three injury models is the associated disruption of the blood-brain barrier (BBB), which could result in the elevation of various cytokines and inflammatory factors at the lesion site. The endogenous, CNS-based expression of cytokine mRNAs was tested by qPCR at different time points after the injury, with GFAP gene activity as internal indicator of glial activation (**Supplementary Fig. 10a**). And indeed, the endogenous expression of interleukin-6 (IL-6), bone morphogenetic protein 4 (BMP4), leukemia inhibitory factor (LIF) and ciliary neurotrophic factor (CNTF) was significantly up-regulated after SWI, however at different time scales (**Supplementary Fig. 10b-e**).

**Figure 6.**
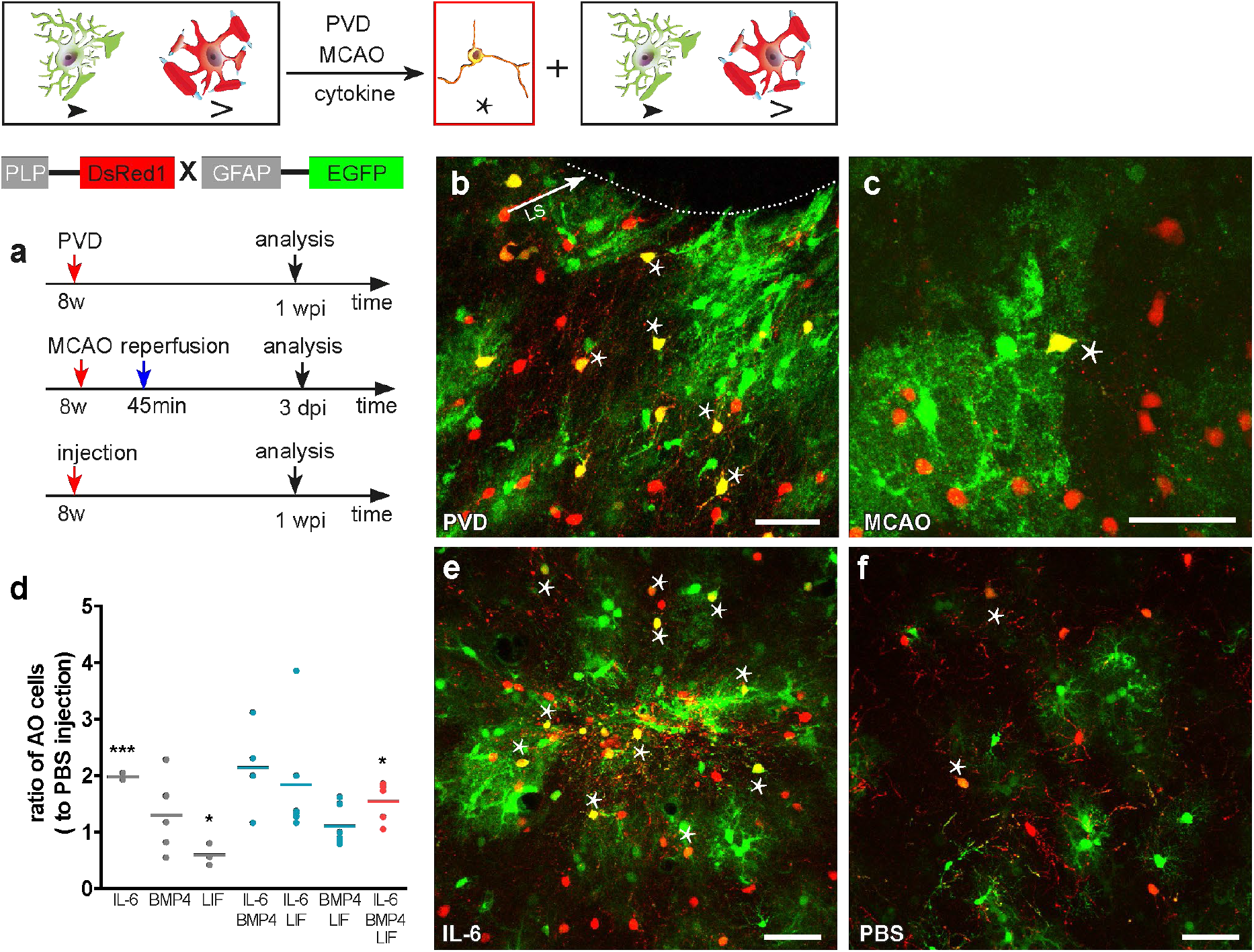
Cytokines modulate AO cell activation after SWI. **a**, Experimental schedules. **b, c**, AO cells (asterisks) appeared adjacent to the lesion (LS) one week after PVD (arrow, **b**) and in penumbra three days after MCAO (45 min occlusion, **c**). **d-f**, AO cells formed at 1 w after BMP4, IL-6 (**e**), LIF, BMP4/IL-6, BMP4/LIF, IL-6/LIF and BMP4/IL-6/LIF injection. Quantification of relative AO cell numbers revealed distinct impact of cytokines on oligodendroglial plasticity, compared with PBS injection (**d**). ^*^: p < 0.05, ^***^: p < 0.001, compared with PBS injected site, one-way ANOVA. Asterisks indicate AO cells. Scale bars in **b, c** = 25 µm; **e, f** = 50 µm.

BMP4 and IL-6 induce neural stem cells and OPCs to differentiate into astrocytes rather than oligodendrocytes^19, 20^, while LIF facilitates oligodendrocyte differentiation^21^. To investigate whether these cytokines could regulate the fate decision of oligodendrocytes *in vivo*, we investigated the number of AO cells in PLP-DsRed1/GFAP-EGFP mice (**Fig. 6d-f**) after cortical cytokine injection. AO cell numbers increased significantly after IL-6 injection (**Fig. 6d, e**, 1.98 ± 0.04 fold, n = 3 mice, p = 1.37E-05), but not after BMP4 injection (**Fig. 6d**, 1.3 ± 0.3 fold, n = 5 mice) compared to PBS injection (**Fig. 6f**). In contrast, LIF reduced AO cell numbers (**Fig. 6d**, 0.6 ± 0.11 fold, n = 3 mice, p=0.0227) compared to PBS. Our data suggest IL-6 as a predominant factor of differentiation, since also the combination IL-6/BMP4/LIF (**Fig. 6d**, 1.4 ± 0.18, n = 6, p = 0.0435) increased the relative quantity of AO cells, despite the reductive effect of LIF. In conclusion, our data suggest that AO cell formation is promoted by IL-6 and inhibited by LIF.

## Discussion

In this study, we report that the damage associated with an acute brain trauma triggers oligodendrocytes to acquire a bi-potential cell lineage commitment via a transitional cell stage. This transitional AO cells (termed according to their <u>a</u>stroglial and <u>o</u>ligodendroglial properties) can change their oligodendroglial lineage fate and become astrocytes. The differentiation of AO cells depends on cortical niches and is influenced by cytokines in the acute lesioned area (**Fig. 7**).

**Figure 7.**
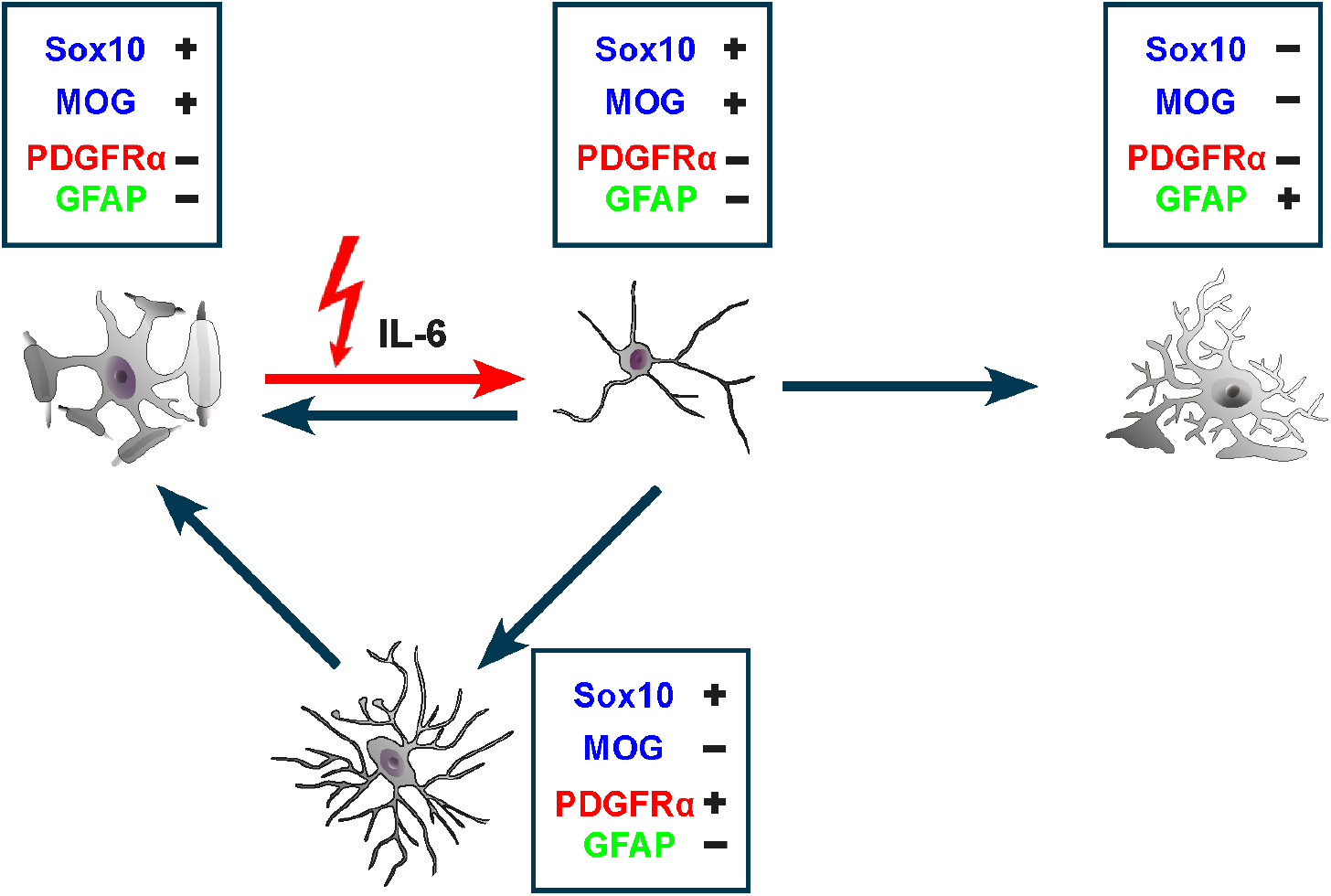
Oligodendroglial fate plasticity induced by acute cortical injuries. A subpopulation of oligodendrocytes (So×10^+^, MOG^+^, PLP^+^, MAG^+^, PDGFRα^-^, GFAP^-^) activate the GFAP promoter induced by acute cortical injuries (SWI, PVD and MCAO) via IL-6, enter a transitional AO cell stage (So×10^+^, MOG^+^, PDGFRα^-^, GFAP^-^) and further give rise to astrocytes (So×10^−^, MOG^-^, PDGFRα^-^, GFAP^+^, GS), oligodendrocytes (So×10^+^, MOG^+^, PDGFRα^-^, GFAP^-^) or OPCs (So×10^+^, MOG^-^, PDGFRα^+^, GFAP^-^).

### Oligodendrocytes regain a plastic phenotype after acute brain injuries

Oligodendrocytes are commonly regarded as mature, non-proliferating and terminally differentiated cells. However, over the last decades we could observe accumulating evidence for a more plastic cell fate of oligodendrocytes. Since these data were mainly obtained by studies of lower vertebrates, in human tumor tissue or in the peripheral nervous system^2, 22-24^, their relevance for our understanding of oligodendrocytes in the adult mammalian CNS remained limited. In gold fish, for example, oligodendrocytes dedifferentiate to bipolar cells with retracted myelinating processes upon retinal axon degeneration^3^, morphologically similar to AO cells. But also in mammals, injuries can disconnect myelinating processes from oligodendroglial cell bodies. In *ex vivo* preparations of the mouse optic nerve, oxygen-glucose deprivation (mimicking a stroke injury) induced a loss of myelinated oligodendrocyte processes^25^. In the periphery, myelinating Schwann cells dedifferentiate after nerve injuries. They rapidly down-regulate myelin proteins (peripheral membrane protein 22, myelin basic protein, periaxin) and subsequently express marker proteins for non-myelinating Schwann cells such as GFAP^22, 26^. In humans, some oligodendroglioma cells were found to express GFAP^23, 24^. Such gliofibrillary (GFAP-expressing) oligodendrocytes (GFOCs) were also described as transitional cells exhibiting astrocytic and oligodendroglial properties^27-29^ with the ability to differentiate into astrocytes. The latest evidence was provided by combining MCAO and MBP-Cre/loxP fate mapping. Epigenetic characterization of the *gfap* promoter region in MBP-Cre reporter labeled oligodendrocytes described a putative mechanism how to generate a cell with astroglial properties^6^.

In our study we combined extended neurogenetic approaches and direct *in vivo* visualization of various stages of an injury-evoked conversion of oligodendrocytes to astrocytes and could highlight the enormous plasticity of even terminally differentiated cells dormant in the mammalian CNS.

### AO cells are novel transitional precursors

The morphology of AO cells resembles that of bipolar O-2A progenitor cells displaying round somata and few, fine processes, first described in the 1980s^30^. O-2A cells belong to the oligodendrocyte lineage and give rise to astrocytes and oligodendrocytes *in vitro* dependent on fetal calf serum (FCS) or bone morphogenetic proteins (BMPs)^31, 32^. Similarly, the AO cells described here differentiate also into both cell types, but *in vivo*, and their fate is also modulated by local cues, provided by the complexity of the cortical niche.

While O-2A cells (subsequently characterized as OPCs) express NG2 and PDGFRα^33^, AO cells of our study were always negative for OPC or stem cell markers. Electrophysiological properties of AO cells such as membrane resistance, resting membrane potential and K^+^ current expression were quite variable, thereby stressing a transitional status. This is also indicated by the changing levels of PLP and GFAP promoter activities, i.e. DsRed1/EGFP expression ratios.

The *in vivo* 2P-imaging visualizes that AO cells originate from oligodendrocytes. This observation was further substantiated by the analysis of DsRed1 expression. After prolonged promoter activity and, hence, long-term expression, the reef coral fluorescent protein DsRed1 precipitates in the cytosol and forms fluorescent aggregates. Since we found such DsRed1 deposits in processes of newly generated AO cells of PLP-DsRed1/GFAP-EGFP mice after injury, the DsRed1 expression must have commenced in oligodendrocytes before the injury. The related protein mRFP1 precipitates as well after prolonged expression. Since in PLP-EGFP_mem_/GFAP-mRFP1, we observed a homogenous distribution of mRFP1 (similar to any other soluble protein), this finding implies only a short period of GFAP promoter activity; hence AO cells, formed from green fluorescent (membrane-bound EGFP) oligodendrocytes, activate the GFAP promoter, but express mRFP1 only shortly. This data demonstrate a continuous transformation of one cell type (oligodendrocyte) to another (astrocyte) with AO cells in between. Upon injury AO cells are an additional transitional stage of a glia-restricted precursor cells different to OPCs, representing another cellular component of oligodendroglial heterogeneity^34^.

### Influence of cytokines on AO cell formation and subsequent differentiation fate

In the injured CNS, the levels of cytokines or growth factors are modulated by different mechanisms. Acute brain traumata (SWI, PVD or MCAO) result in BBB disruption and a subsequent influx of different peptides or factors from the peripheral blood into the CNS. But also the activity of endogenous genes contributed to enhanced levels of various cytokines as described previously^35, 36^, and in the current study.

IL-6 can activate the GFAP promoter^37^ as shown by increased AO cell numbers after IL-6 injection. In contrast, LIF^38^) keeps cells in the oligodendrocyte lineage^21, 39^, in line with our observation that LIF injection inhibits the formation of AO cells. However, *in vitro* experiments suggest an astrogliogenic function for LIF^6, 40^. While the acute injection of LIF inhibits AO cell formation, injury evoked LIF might be responsible for astrocyte formation of AO cells. The same might be true for BMP4, which has been suggested to induce astrocytic differentiation from OLCs^41, 42^. Single BMP4 injections did not increase AO cell numbers, in contrast to a combined injection with IL-6 and LIF, which indicates the BMP4 function in inducing astrocyte differentiation, as shown for precursor cells^40^. IL-6, LIF and BMP4 act in a defined temporal pattern of cytokine activity in post-injury processes. IL-6 induces AO cell formation from oligodendrocytes and LIF and BMP4 subsequently affect the astrocyte-specific differentiation. This temporal expression pattern of cytokines is tightly coupled to their local distribution within the cortical layers and in distance to the injury site, thereby forming transient differentiation niches^7, 12, 43–45^.

### Abundance of astrocytes that are derived from oligodendrocytes

Already previous studies suggested that cells of the oligodendrocyte lineage, mainly OPCs, could generate astrocytes after brain damage. But to which extent remained controversial, attributable to the variety of animal models (transgenic mice and rats), lesion paradigms, duration of recombination after induction or other methodologies^9, 11–14, 46^. For example, shorter time intervals left for recombination intervals (like 3 to 5 days after tamoxifen injection) preferentially results in more labelled OPCs rather than oligodendrocytes^8^. In some recent studies, about 5 % and 5.7 % GFAP^+^reporter^+^ astrocytes (of total recombined cells) could be found after SWI^12, 13^. In NG2-CreER^T2^ x R26-tdT mice we observe a very reliable CreERT2 expression controlled from the complete NG2/cspg4 locus^8^. We detected a high percentage of GFAP^+^tdT^+^ cells (25.5 %). Out of these, about 30 % were astrocytes (7.1 % / 25.5 % PDGFRα −) and about 70 % were OPCs (18.4 % /25.5 % PDGFRα ^+^). We observed even more astrocytes (14.1 % vs. 7.1 %) when we analyzed mice with a higher percentage of recombined oligodendrocytes, which duplicate between 10 and 30 d post tamoxifen injection in the cortex of P60 mice^47^. In all of the various mouse lines of our study, we detected AO cells differentiating to astrocytes, however, the quantities of AO cell-derived astrocytes differed due to various degrees of transgenic modifications. In NG2-CreER^T2^ mice all cells of the oligodendrocyte lineage including their putative progeny (AO cells, oligodendrocytes and OPCs) are labeled, while in split-Cre mice (generated by non-homologous recombination) only AO cells and their progeny can be observed. Therefore, the 7.1 % (1 wpi) labeled astrocytes from all recombined cells in NG2-CreER^T2^ mice appear small when compared to split-Cre mice (68.1 %, 1 wpi), but they are calculated from different reference points. Notably, a cell population that increased with time escaped the three-color immunostaining detection. These cells were highly likely non-activated GFAP negative astrocytes frequently observed in the cortex of healthy mice^48^.

### Heterogeneity of astrocytes

During injury-induced proliferation, the number of astrocytes increases by 10 -20 %^49, 50^. For split-Cre mice we estimated the percentage of astrocytes derived from AO cells/oligodendrocytes to reach up to 5 % (tdT^+^GFAP^+^ of total GFAP^+^ cells). Obviously, astrocytes originating from oligodendrocytes constitute a significant portion of cells in the glial scar. Given the limited penetrance of the split-Cre transgenes, this number could even be higher^51^. Whether these astrocytes fulfill a specialized function during scar formation or perform more classical astroglial tasks such as regulating extracellular ion and transmitter homeostasis or enhancing neuronal energy supply remains to be determined. Very recently, two different classes of astrocytes were described within a glial scar. While A1 astrocytes were triggered by microglial cytokine release and became neurotoxic, A2 astrocytes started to release various growth factors and appeared to be neuroprotective^52, 53^. Remembering the early *in vitro* work of Ffrench-Constant and Raff, it is very tempting to speculate that AO cell-derived astrocytes could comprise a major portion of the A2 astrocytes^54^. Local, environmental cues and controlled switching of cell lineages generate a novel dimension of heterogeneity^55^. And indeed, astrocytes display unique profiles of gene expression and cell behavior after acute and chronic injuries^56^.

Targeting the plasticity of oligodendrocytes as well as the function of oligodendrocyte-derived astrocytes could become an exciting field to explore novel routes in treating acute brain trauma.

## Materials and Methods

### Ethics statements

This study was carried out at the University of Saarland in strict accordance with recommendations of European and German guidelines for the welfare of experimental animals. Animal experiments were approved by Saarland state’s “Landesamt für Gesundheit und Verbraucherschutz” in Saarbrücken/Germany (animal license numbers: 71/2010, 72/2010 and 65/2013, 36/2016, 03/2021).

### Animals

Mouse breeding and animal experiments were performed in the animal facilities of the University of Saarland. In this study, heterozygous male and female mice at the age of 8-14 weeks were used, except that P20 mice for electrophysiological studies. Split-Cre DNA recombinase mice for coincidence detection as well as inducible Cre DNA recombinase mice (NG2-CreER^T2^, GFAP-CreER^T2^, PLP-CreER^T2^ and Glast-CreER^T2^) were always used in combination with floxed reporter (homozygous for Rosa26-EYFP or Rosa26-tdTomato, **Supplementary Table 2**) mice to show successful recombination. Different transgenic split-Cre mice were analyzed and the double transgenic line showing the lowest cortical recombination after embryonic development was used (GCNT x PCCK). In addition, we crossed the NG2-CreER^T2^ mice with green astrocyte specific fluorescent mice GFAP-EGFP_GFEC_(later referred to as NG2-CreER^T2^ x GFAP-EGFP). Fluorescent astrocyte mouse line (GFAP-EGFP_GFEA_) was bred to an oligodendroglial specific mouse with DsRed1 expression under control of the murine PLP promoter, and a GFAP-mRFP1_GRFT_ mouse line to PLP-EGFP_mem_. A more detailed description of tamoxifen protocols and mouse lines can be found in **Supplementary Table 2 online**.

### Acute brain injury models

Stab wound injury (SWI) was performed in young (P20) or adult anesthetized mice (ketamine (Ketavet®, Pfizer, Germany) / xylazine (Rumpon®, Bayer Healthcare, Germany) in 0.9 % NaCl (140 mg/10 mg per 1 kg body weight)). The skull was thinned with dental drill laterally 1.5 mm and longitudinally 2 mm from bregma, followed by a 1 mm deep stab wound made with a scalpel.

For pial vessel disruption (PVD) a cortical craniotomy (3 mm diameter, center: approximately located (bregma considered as 0) laterally 1.5 mm and longitudinally 2 mm) was made in anesthetized mice^57^. A medium (class II) vessel was disrupted with sharp forceps (#5, Fine Science Tool, Heidelberg, Germany) without interference of larger (class I) vessels. Bleeding was stopped with ice-cold 0.9 % NaCl solution.

Middle cerebral artery occlusion (MCAO) was performed as described with few modifications^58^. Briefly, after 45 min of occlusion, the filament was removed and the wound closed. For energy recovery, 10 % glucose (1 ml/20 g body weight) solution was injected i.p.

Cytokine injection was made in the center of a cortical craniotomy. A glass pipet was filled with 0.3 µl of PBS, BMP4 (500 µg/ml in PBS, Abcam, Cambridge, UK), LIF (200 µg/ml in PBS, Isokine, Kopavogur, Iceland), IL-6 (100 µg/ml, Biomol, Hamburg, Germany) or a combination of BMP4/LIF, BMP4/IL-6, IL-6/LIF or BMP4/IL-6/LIF. Injection was performed with a programmable syringe pump (540060, TSE systems, Germany).

After surgeries (SWI, PVD, MCAO and cytokine injections) the wound was closed and buprenorphine (0.01 µg / 30 g body weight, Temgesic, Essex Pharma, Muenchen, Germany) was injected as anti-pain treatment.

### *In vivo* two-photon laser-scanning microscopy (2P-LSM)

For 2P-LSM a 3 mm-diameter cortical craniotomy (lateral=1.5 mm and longitudinal=2 mm from bregma) was made^59^. The center was stabbed with a needle (0.46 mm x 2 mm) and rinsed with cortex buffer (in mM: 125 NaCl, 5 KCl, 10 glucose, 10 HEPES, 2 CaCl_2,_ 2 MgSO_4_ (pH∼ 7.4)) until the bleeding stopped. A 3 mm coverslip was placed on the brain and fixed with dental cement (RelyX^®^, 3M-ESPE, Neuss, Germany). More than twenty GFAP-EGFP/PLP-DsRed1 and ten PLP-EGFP_mem_/GFAP-mRFP1 mice were investigated up to ten times in between 30 days after the injury at different time intervals. However, a several AO cells could not be followed due to technical problems such as window blurring upon glial scar formation or bone regrowth. To be as stringent as possible, we disregarded all observations on “AO cells becoming astrocytes” when we could not unequivocally identify the oligodendroglial origin of an AO cell, even when thereby sacrificing the percentage of astrocytes coming from an oligodendrocyte.

### Immunohistochemistry

Mouse perfusion, tissue fixation and vibratome slice preparation (40 µm) was performed as described previously^8^.

For fluorescent immunohistochemistry, the following primary antibodies were used: polyclonal goat: anti-GFP (1:1000, Rockland, Gilbertsville, USA)) and anti-PDGFRα (1:500, R&D Systems, Minneapolis, USA), anti-Sox10 (1:100, R&D Systems, Minneapolis, USA); polyclonal rabbit: anti-GFAP (1:1000, Dako Cytomation, Glostrup, Denmark), anti-Cre (1:500, Novagen, Darmstadt, Germany), anti-S100B (1:500, Abcam, Cambridge, UK), anti-Iba1 (1:1000, Wako, Neuss, Germany), anti-Sox2 (1:200, R&D Systems, Minneapolis, USA), anti-DsRed (1:1000, Clontech, Mountain View, USA) and anti-Olig2 (1:200, Dr. C. Stiles, Harvard Medical School); monoclonal mouse: anti-GSTπ (1:500, BD Transduction Laboratories, San Jose, USA), anti-glutamine synthethase (GS) (1:500, Transduction Laboratories, San Jose, USA), anti-MAG (1:500, Merck Millipore, Darmstadt, Germany), anti-MOG (1:500, Abcam, Cambridge, UK), and anti-Camsap1 (1:500, Dr. Hiroaki Asou), anti-PLP (1:500, Dr. K. Nave), anti-NeuN (1:500, Merck Millipore, Darmstadt, Germany); monoclonal rat anti-NG2 (1:200, Dr. J. Trotter), anti-BrdU (1:1000, Abcam, Cambridge, UK). Secondary antibodies were Alexa488/633-conjugated anti-mouse, Alexa555-conjugated anti-rabbit, Alexa488/633-conjugated anti-goat (all 1:2000, Invitrogen, Grand Island NY, USA) and Cy5 anti-rat IgG. Please note, Cre immunostaining recognized only the larger C-terminal, but not the shorter N-terminal fragment of the Cre DNA recombinase, therefore only PLP-promoter active cells could be labelled.

For DAB staining, after primary antibody incubation, biotinylated secondary antibody was incubated, followed by incubation with freshly prepared AB mix of Vector Elite ABC Kits and with DAB (DAKO Agilent Pathology Solutions, Santa Clara, CA, USA).

### Fluorescence *in situ* hybridization (FISH)/Immuno-FISH

We performed fluorescence *in situ* hybridization (FISH)/immuno-FISH using the QuantiGene®ViewRNA ISH Cell Assay Kit (Affymetrix) with few modifications from the manual. Briefly, vibratome slices were permeabilized at 4°C for 1 h and washed once with PBS. Then slices were incubated in 40 °C with *GFAP mRNA* probes (1:100, 3 h), pre-amplification solution (1:25, 1 h), amplification solution (1:25, 1 h) and the fluorescent label probe solution (1:25, 1 h) with intermediate washing steps (wash buffer). Finally, the slices are washed once with PBS and can be either directly mounted or stored in PBS at 4°C overnight for further immunohistochemical staining.

### Proliferation analysis

Split-Cre mice received drinking water containing 5-bromo-2’-deoxyuridine (BrdU) (Sigma-Aldrich) (1 mg/ml) for two consecutive weeks following SWI *ad libitum*.

### Tamoxifen-induced gene recombination

To induce reporter expression in CreER^T2^-mice, tamoxifen dissolved in cornoil (10 mg/ml) was intraperitoneally injected (100 mg / kg body weight) to 7-week-old mice once per day for three consecutive days.

### Whole-cell patch-clamp analysis

The lesioned brain of young mice (P20, 3 or 4 dpi) was dissected and placed in ice-cooled, carbogen-saturated Ca^2+^-free preparation solution (in mM: 126 NaCl, 3 KCl, 25 NaHCO_3_, 1.2 NaH_2_PO_4_, 3 MgCl_2_ and 15 Glucose). Acute 300 μm frontal vibratome sections (Leica VT 1200S) were obtained. After at least 1 h recovery in oxygenated aCSF (in mM: 126 NaCl, 3 KCl, 25 NaHCO_3_, 15 glucose, 1.2 NaH_2_PO_4_, 2 CaCl_2_, and 2 MgCl_2_ at 35°C), they were subsequently transferred to a recording chamber mounted on an upright microscope (Axioscope 2 FSmot, Zeiss, Germany) and continuously perfused with aCSF (126 NaCl, 3 KCl, 25 NaHCO_3_, 15 glucose, 1.2 NaH_2_PO_4_, 1 MgCl_2_and 2.5 CaCl_2_, room temperature; 20–23°C) at a flow rate of 2–5 ml/min. AO cells, astrocytes, oligodendrocytes and OPCs were identified by their respective fluorescence using conventional epifluorescence illumination. Images were taken with a CCD camera (Kappa, DX4C-285FW). Whole-cell voltage-clamp recordings were obtained with an EPC10 patch-clamp amplifier (HEKA, Lambrecht/Pfalz, Germany), low pass-filtered at 3 kHz and data acquisition was controlled by Patchmaster (HEKA). Currents were recorded at 20 kHz. Patch electrodes were pulled from borosilicate glass capillaries (OD: 1.5 mm; Hilgenberg GmbH, Germany) using a micropipette puller (Model P-97, Sutter Instruments Co., CA) and had a resistance between 4 and 7 MΩ. Patch pipettes were filled with an intracellular solution (in mM: 120 KCl, 5 MgCl_2_, 5 EGTA, 10 HEPES and 5 Na_2_ATP (pH∼ 7.2)).

Analysis was performed with IGOR Pro Version 6.22 (Wavemetrics, Inc., USA), Microsoft Excel and GraphPad Prism 8.0. Glial cells were voltage-clamped at −80 mV (*V*_hold_). Whole-cell membrane currents were evoked by a series of hyper-and depolarizing voltage steps ranging from −140 to 80 mV with an increment of 10 mV.

### Quantitative real-time PCR (qRT-PCR)

Mice (n = 3 for each group) were perfused with PBS at 0.5, 1, 3, 5 and 7 dpi. Cortical mRNA from ipsi-and contralateral tissue was collected within 0.2 mm thickness and 2 mm width (1 mm from lesion site to each direction). The level of mRNA was detected by qRT-PCR. Primers for qRT-PCR were as follows (in 5’ to 3’ direction): BMP4-forward, GAG CCA TTC CGT AGT GCC AT; BMP4-reverse, ACG ACC ATC AGC ATT CGG TT; IL-6-forward, GAG TGG CTA AGG ACC AAG ACC; IL-6-reverse, AAC GCA CTA GGT TTG CCG A; LIF-forward, CCC AGC ATC CCA GAA CCA TT; LIF-reverse, AGA GCT GGG TTG CTT GAG TC; GFAP-forward, TGG AGG AGG AGA TCC AGT TC; GFAP-reverse, AGC TGC TCC CGG AGT TCT; CNTF-forward, GAC CTG ACT GCT CTT ATG GAA TCT; CNTF-reverse, AGG TTC TCT TGG AGG TCC G; ATPase-forward, GGA TCT GCT GGC CCC ATA C; ATPase-reverse, CTT TCC AAC GCC AGC ACC T.

### Microscopic analysis and quantification

Three brain slices per mouse and at least three animals per group were examined. Overview images were obtained with an epifluorescence microscope (Eclipse E600, Nikon, Japan) using NIS-Elements BR (Nikon version 2.2). Confocal images were taken by a laser-scanning microscope (LSM-710, Zeiss), processed with ZEN software (Zeiss) and displayed as single optical sections, orthogonal image stacks or maximum intensity projections. For statistical analysis raw or linearly processed image data were used. Figures presented in this work were modified with image processing tools of ImageJ (Fiji, www.fiji.sc) and Zen 2011 software (Zeiss, Oberkochen, Germany). For cell counting in NG2-CreER^T2^ and NG2-CreER^T2^ x GFAP-EGFP mice the also NG2-positive, vessel-associated pericytes were excluded by their morphology. For region-dependent cell counting, 6-8 z-stacks were taken without overlap along the lesion site and three stacks at the contralateral side. The volumes for cell counting are listed in **Supplementary Table 3 online**. We performed double-immunostainings to identify the glial cell types that contribute to the recombined (tdT+) cell population (**Supplementary Fig. 4**). Please note that only three detection channels were available for cell characterization (tdT+ and two for GFAP, PDGFRα or GSTπ). Therefore, we always were left with an unidentified cell population (**Supplementary Fig. 4b-e**). Since this population increased with time after injury, they probably were astrocytes that had down-regulated their GFAP expression, a phenomenon common to non-activated cortical astrocytes.

In all figures the following symbols were used for the different cell types: triangles indicate OPCs, open triangles oligodendrocytes, arrowheads astrocytes and asterisks AO cells.

## Statistical analysis

Three animals of every experimental age group and mouse line were studied in three independent experiments. Cells counted at the lesion site were observed in a region 0-300 µm from the lesion in both directions. Statistical differences were analyzed using the two-tailed *t*-test for two-group comparison, one-way ANOVA for comparison among more than two groups. Data are shown as mean ± SEM.

## Data availability

All data, materials and protocols are available from the corresponding authors on reasonable request.

## Supplementary Figure legends

**Supplementary Figure 1.**
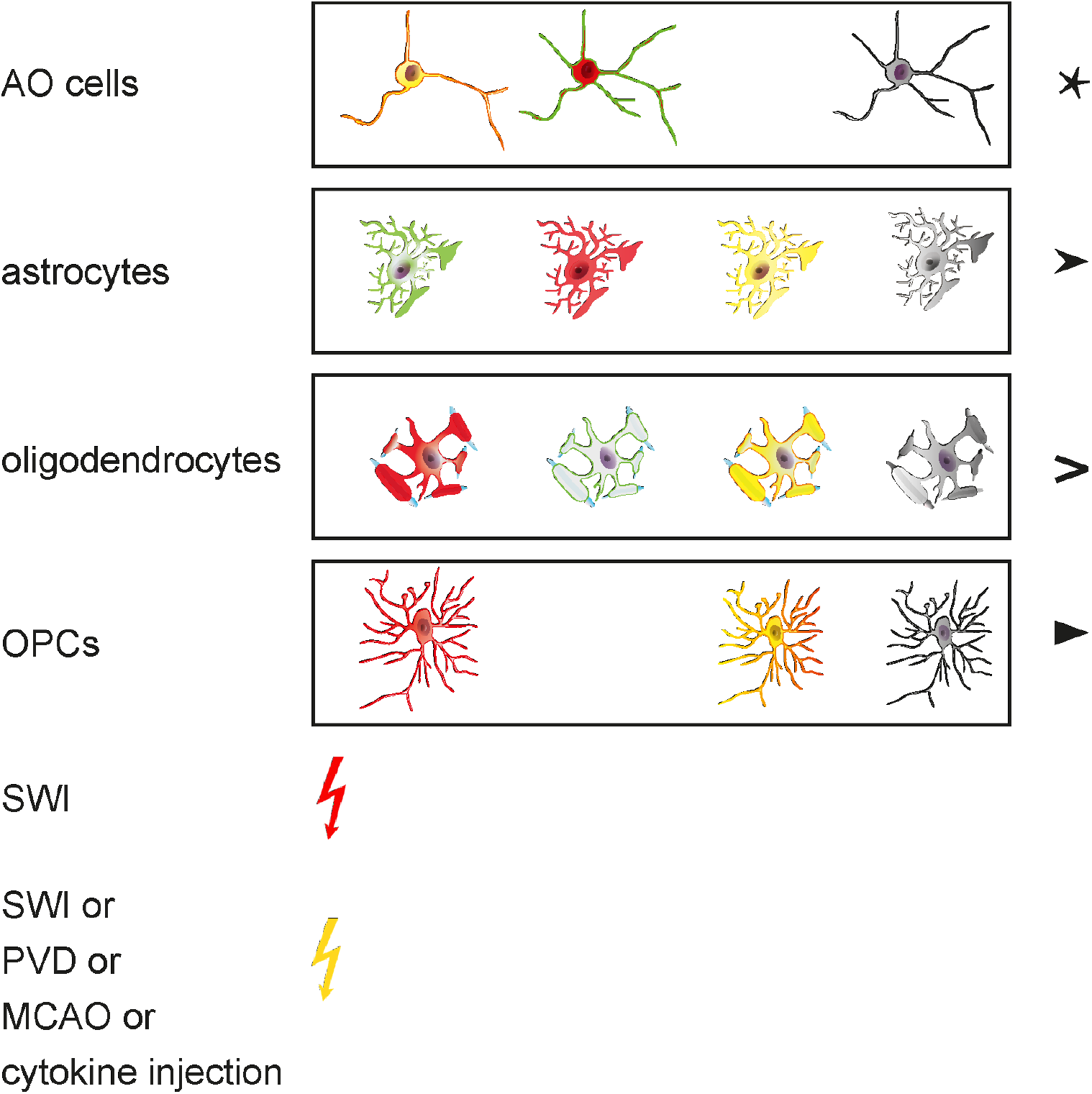
Symbols for glial cells indicating transgenic protein expression and injury models as used in the figures.

**Supplementary Figure 2.**
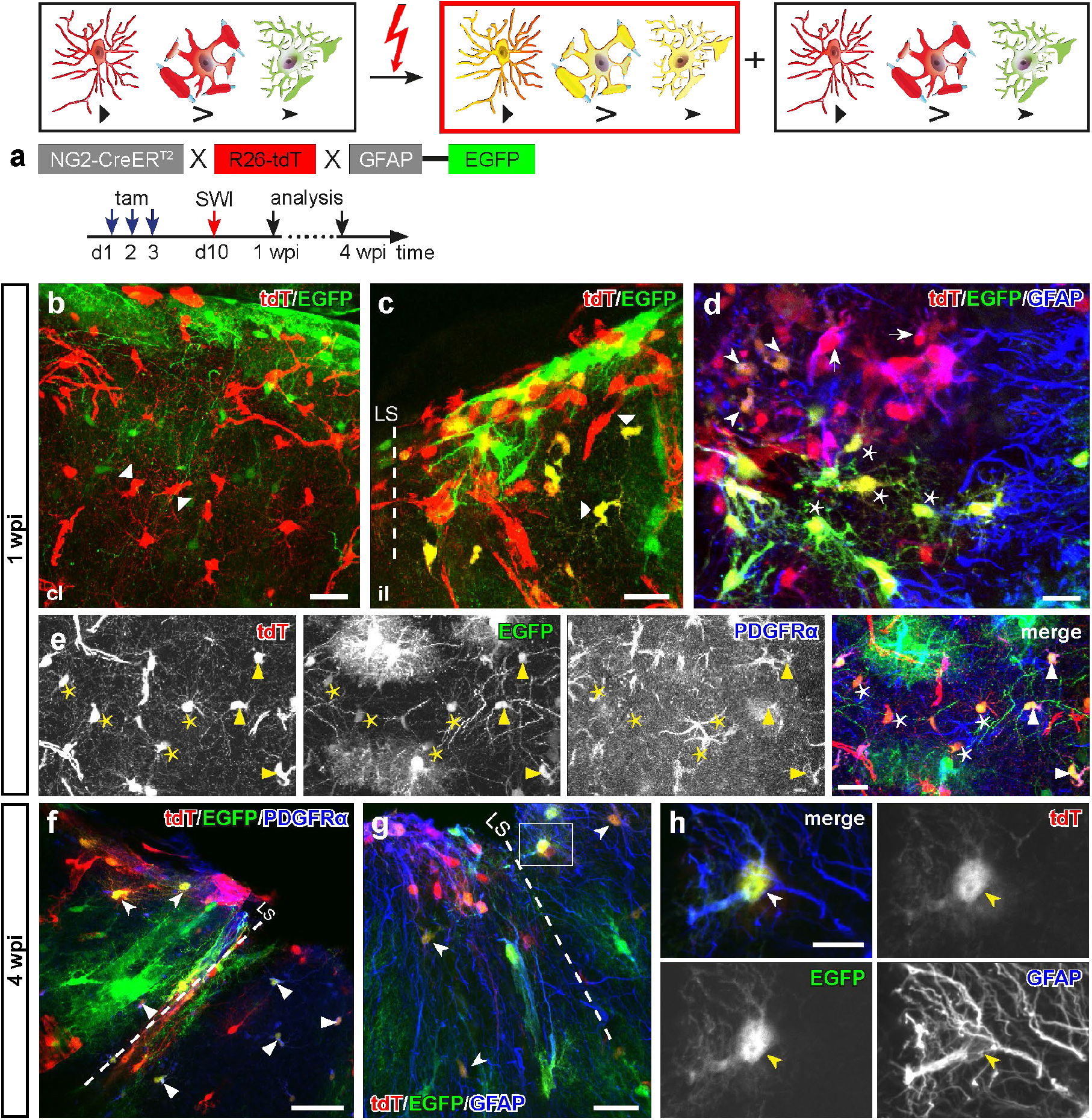
Mature oligodendrocytes can activate the transgenic GFAP promoter. Crossbreeding of GFAP-EGFP_GFEC_ to NG2-CreER^T2^ x R26-tdT mice facilitated the observation of OPCs (triangles) and AO cells (asterisks) possessing transgenic GFAP promoter activity. **a**, Protocol of DNA recombination induction and analysis of NG2-CreER^T2^ x GFAP-EGFP_GFEC_ mice. **b**, No overlay of EGFP and tdT in the contralateral side. **c**, At the lesion site, a high quantity of cells co-expressed tdT and EGFP 1 wpi, some with the morphology of NG2 glia (triangle). **d**, Two subpopulations of tdT^+^EGFP^+^ cells could be found: tdT^+^EGFP^+^GFAP^+^ (arrowheads, regarded as astrocytes) and tdT^+^EGFP^+^GFAP^-^ (asterisks, AO cells). Since in the transgenic GFAP-EGFP_GFEC_ mice only 38.5 ± 6.3 % of S100B+ astrocytes expressed EGFP in the cortical grey matter, tdT^+^GFAP^+^EGFP^-^ astrocytes were found as well (arrow). **e**, Observation of EGFP^+^ OPCs (EGFP^+^tdT^+^PDGFRα^+^, triangles, 51.6 ± 2.5 %, 97/194 cells, n = 3) and oligodendroglial cells (EGFP^+^tdT^+^PDGFRα^-^, asterisks, 48.4 ± 3.0 %) with transgenic GFAP promoter activity at the ipsilateral side of NG2-CreER^T2^ x GFAP-EGFP mice. **f**, At 4 wpi recombined cells with tdT and EGFP expression can be found either expressing (OPCs, triangles) or lacking PDGFRα (astrocytes, arrowheads). **g**, EGFP^+^tdT^+^GFAP^+^ *bona fide* astrocytes were detectable 4 wpi. **h**, Magnification of GFAP^+^ recombined astrocyte with EGFP and tdT expression 4 wpi. Triangles: OPCs, arrowheads/arrow: astrocytes, asterisks: AO cells. Scale bars in **b-g** = 25 µm, **h**= 10 µm.

**Supplementary Figure 3.**
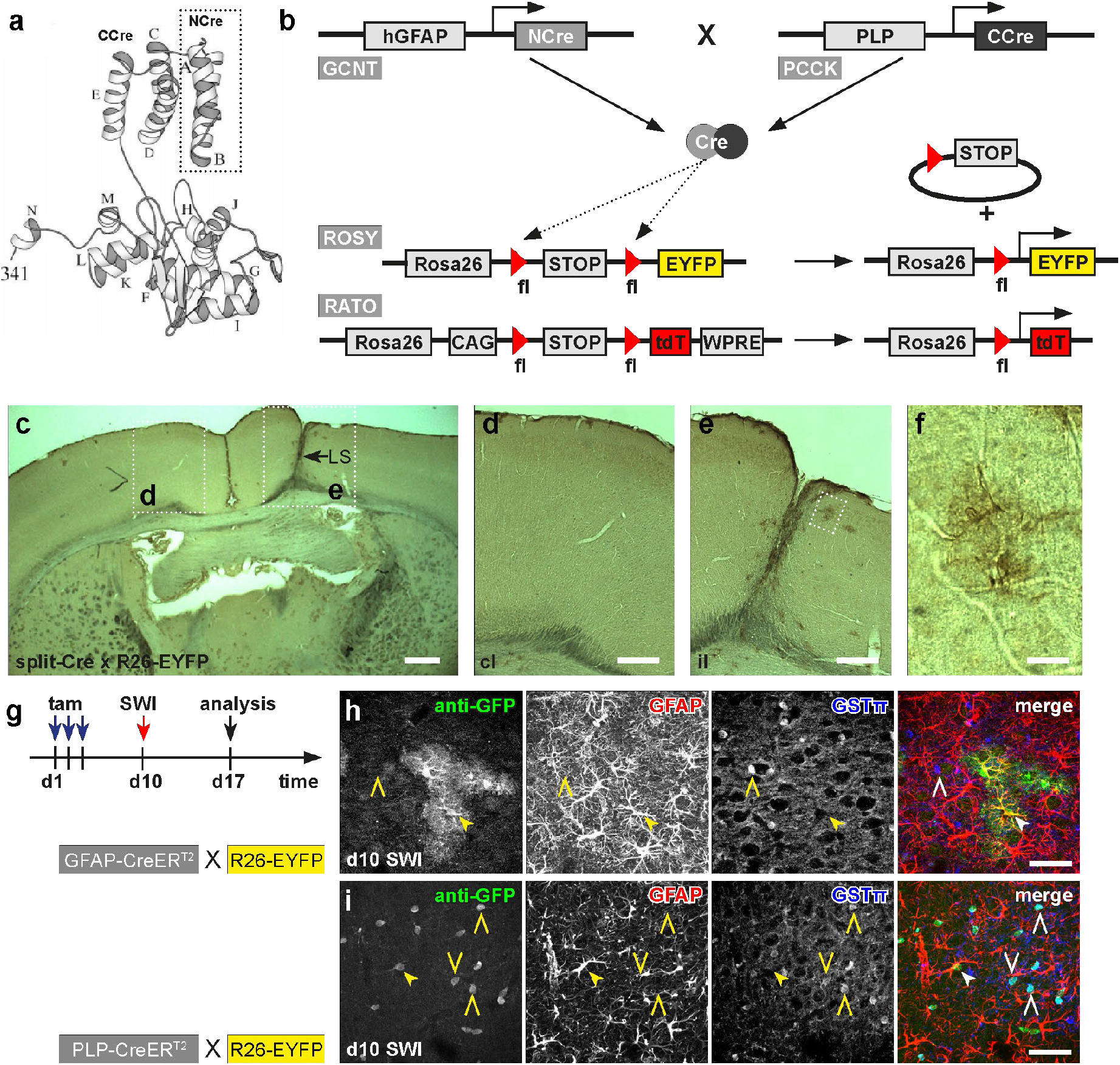
Cortical SWI induces coincident activation of GFAP and PLP promoters. **a**, Split-Cre transgene. **b**, Scheme of split-Cre recombination. **c-f**, DAB staining of GFP in split-Cre x R26-EYFP (Rosa26-STOP-EYFP) mouse 1 wpi revealed exclusive recombination in the lesion site (**c, e**) but not in the contralateral side (**c, d**). Magnified view showed clear morphology of recombined *bona fide* astrocyte (**f**). **g**, Experimental schedules for tamoxifen induced recombination before SWI. **h**, In GFAP-CreER^T2^ x R26-EYFP mice recombined cells were GFAP+ astrocytes, but no oligodendrocytes could be observed. (**i**) In PLP-CreER^T2^ x R26-EYFP mice recombined astrocytes could be detected when the SWI was performed ten days after tamoxifen injection in addition to GSTπ+ oligodendrocytes. Therefore, the transgenic GFAP promoter was activated in recombined oligodendrocytes (PLP-CreER^T2^), while recombined astrocytes (GFAP-CreER^T2^) stay in their lineage. Arrowheads: astrocytes, open triangles: oligodendrocytes. Scale bars in **c** = 500 µm, **d, e** = 200 µm, **f** = 40, **h, i** = 50 µm.

**Supplementary Figure 4.**
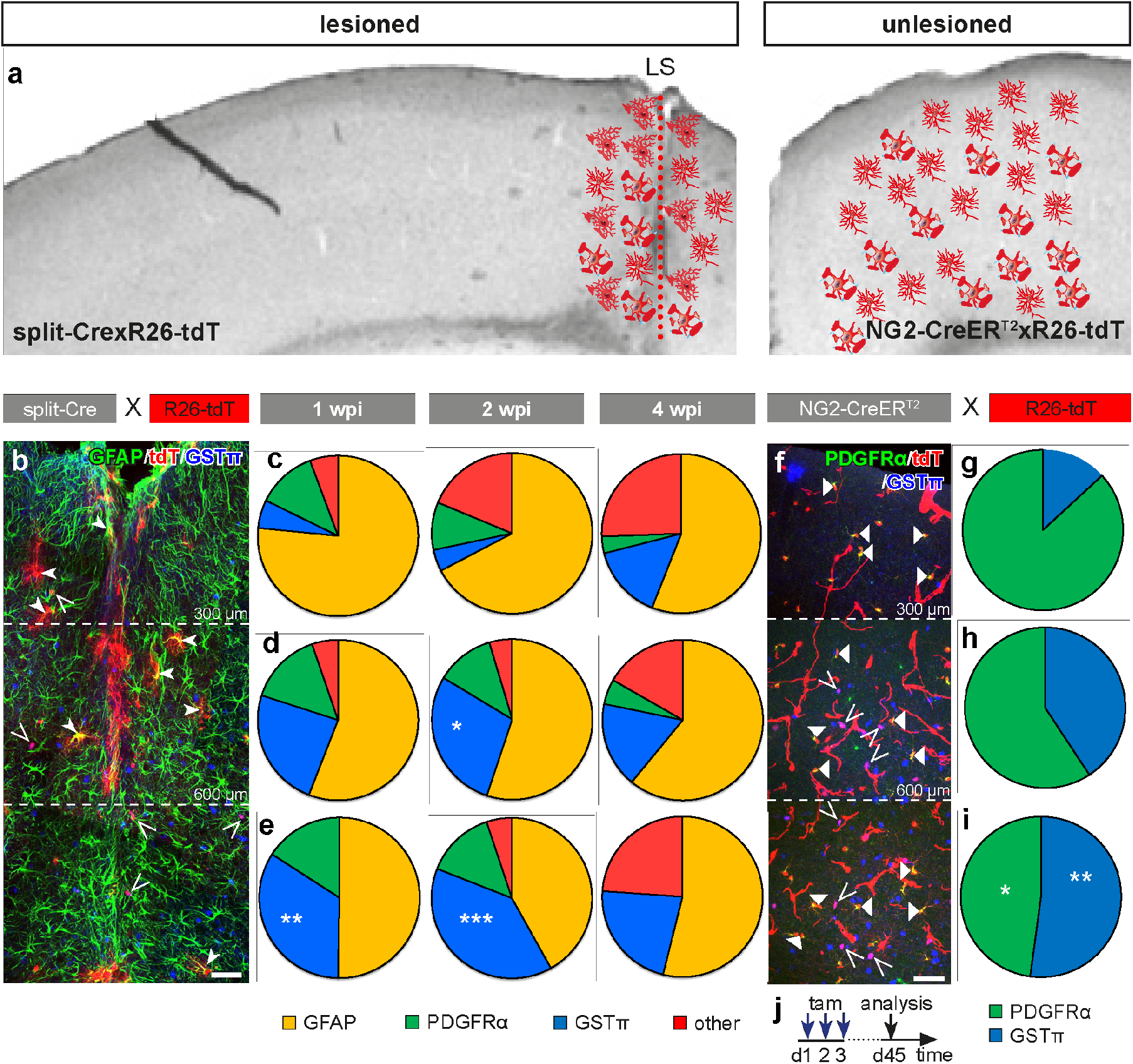
AO cell differentiation depends on local cues. **a**, Scheme of cell distribution at the ipsilateral cortex of split-Cre mice and the intact cortex of NG2-CreER^T2^ mice. **b**, Distribution of recombined cells along the lesion site in split-Cre mice. **c-e**, Quantification of recombined astrocytes (GFAP^+^, yellow), oligodendrocytes (GSTπ^+^, blue), OPCs (PDGFRα^+^, green) and other cells (unidentified, red) in layer I-III (**c**), layer IV-V (**d**) and layer VI (**e**) of split-Cre mice at 1, 2 and 4 wpi. **f-i**, Overview (**f**) and quantification (**g-i**) of the distribution of OPCs (PDGFRα^+^) and oligodendrocytes (GSTπ^+^) in layer I-III (**g**), layer IV-V (**h**) and layer VI (**i**) of the intact cortex of NG2-CreER^T2^ x R26-tdT mice. **j**, Experimental schedule for the analysis of NG2-CreER^T2^ mice. ^*^: p < 0.05, ^**^: p < 0.01 and ^***^: p < 0.001, compared with same population at the same time point to layer I-III, one-way ANOVA. Triangles: OPCs, open triangles: oligodendrocytes, arrowheads: astrocytes. Scale bars in **b, f** = 50 µm.

**Supplementary Figure 5.**
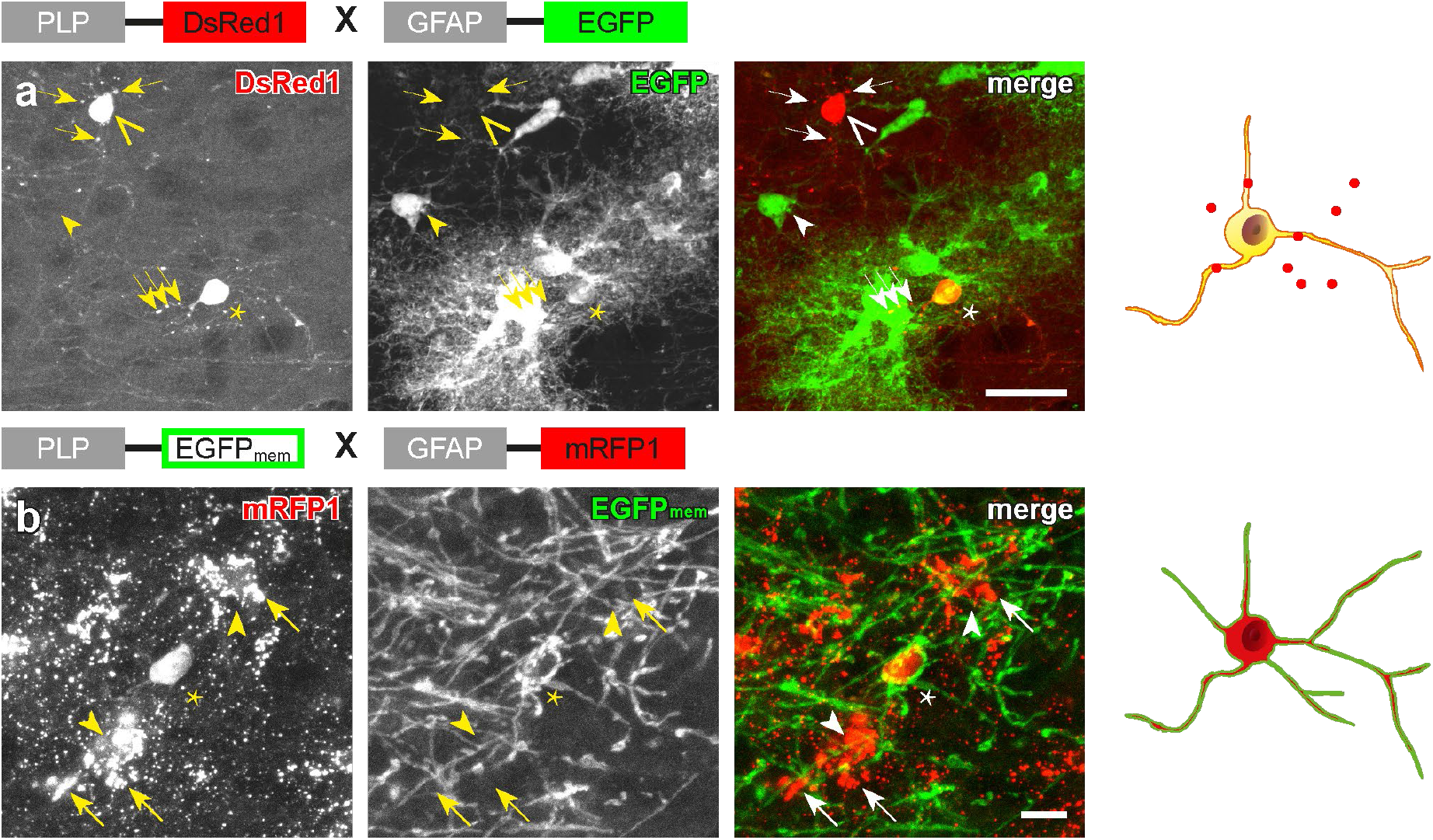
Morphological comparison of AO cells in PLP-DsRed1/GFAP-EGFP and PLP-EGFP_mem_/GFAP-mRFP1 mice. **a**, AO cells in PLP-DsRed1/GFAP-EGFP mouse show DsRed1 clusters (arrows) in the processes, as oligodendrocytes do (open triangle). **b**, AO cells in PLP-EGFP_mem_/GFAP-mRFP1 mouse express cytosolic mRFP1, while surrounding astrocytes display mRFP1 clusters. open triangles: oligodendrocytes, arrowheads: astrocytes, asterisks: AO cells. Scale bars in **a** = 25 µm, **b** = 10 µm.

**Supplementary Figure 6.**
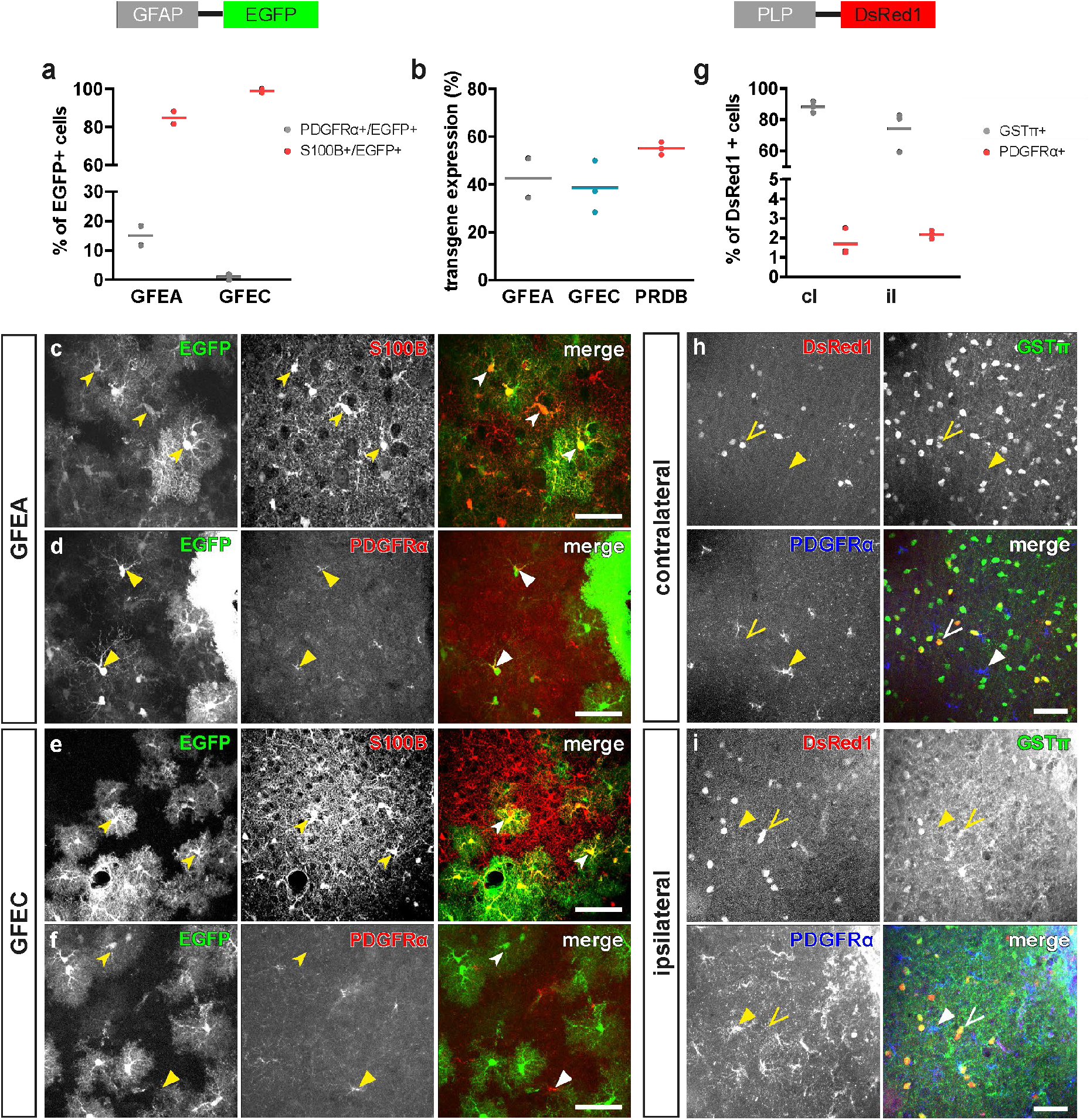
Analysis of transgene expression in both hGFAP-EGFP and PLP-DsRed1 mouse lines. **a-f**, Quantification of EGFP-expressing astrocytes (EGFP^+^S100B^+^) and OPCs (EGFP^+^PDGFRα^+^) cells in both transgenic mouse lines with EGFP expression under control of the human GFAP promoter (hGFAP-EGFP_GFEA_ and hGFAP-EGFP_GFEC_). **b**, Transgene efficacy in hGFAP-EGFP_GFEA_, hGFAP-EGFP_GFEC_ and PLP-DsRed1 mice. **g-i**, Quantification of oligodendrocytes (GSTπ^+^) and OPCs (PDGFRα^+^) expressing DsRed1 in contralateral (**h**) and ipsilateral (**i**) sides of PLP-DsRed1 mouse cortex. open triangles: oligodendrocytes, arrowheads: astrocytes, triangles: OPCs. Scale bars = 50 µm.

**Supplementary Figure 7.**
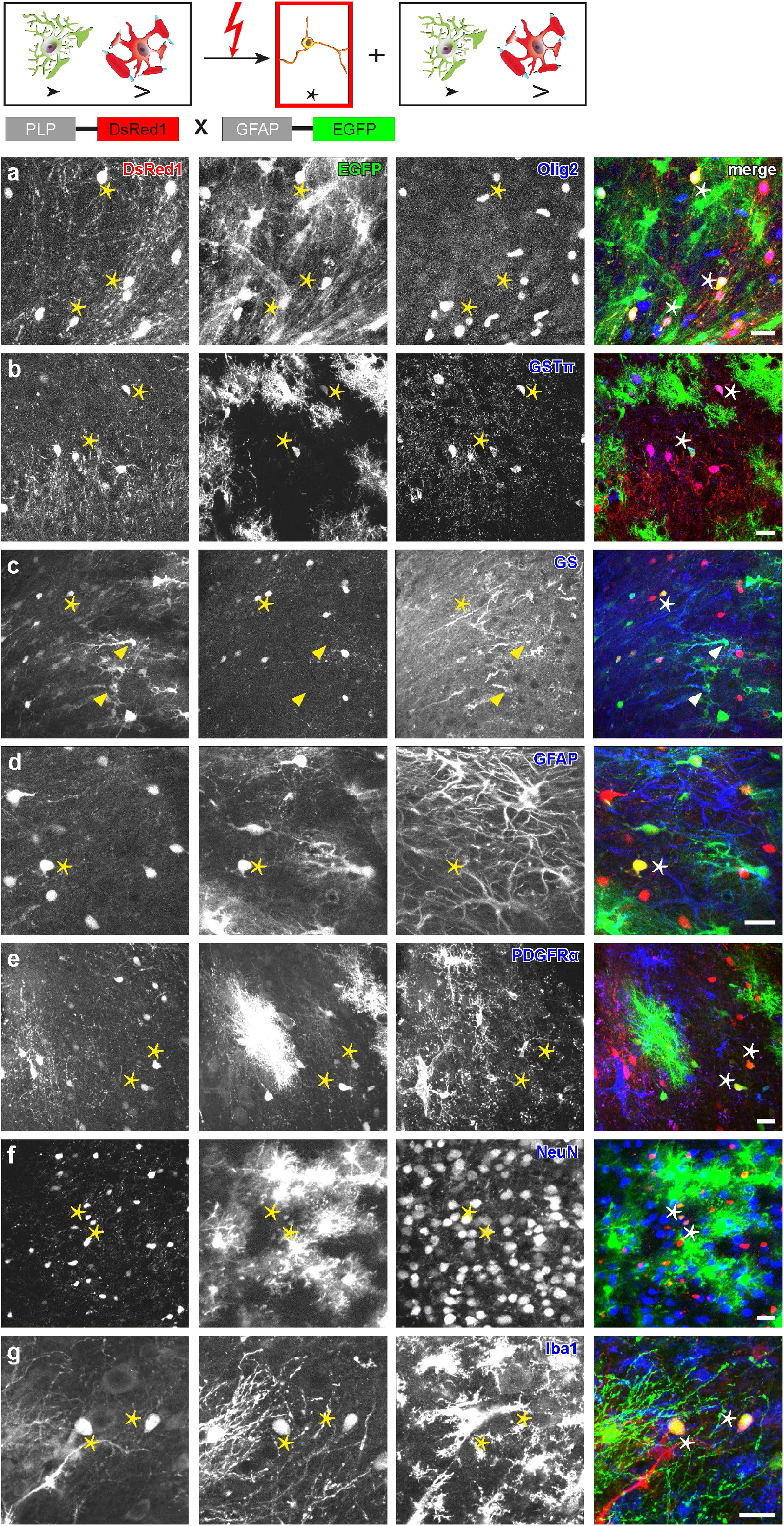
Immunohistochemical analysis of AO cells with cell-specific markers in PLP-DsRed1/GFAP-EGFP mice. **a**, AO cells (asterisks) were positive for the oligodendrocyte lineage marker Olig2. **b, c**, Few AO cells were positive for mature oligodendrocyte marker GSTπ or astrocyte marker GS. **d-g**, AO cells (asterisks) were negative for astroglial (GFAP, **d**), OPC (PDGFRα, **e**), neuronal (NeuN, **f**) and microglial markers (Iba1, **g**). Asterisks: AO cells. Scale bars = 20 µm.

**Supplementary Figure 8.**
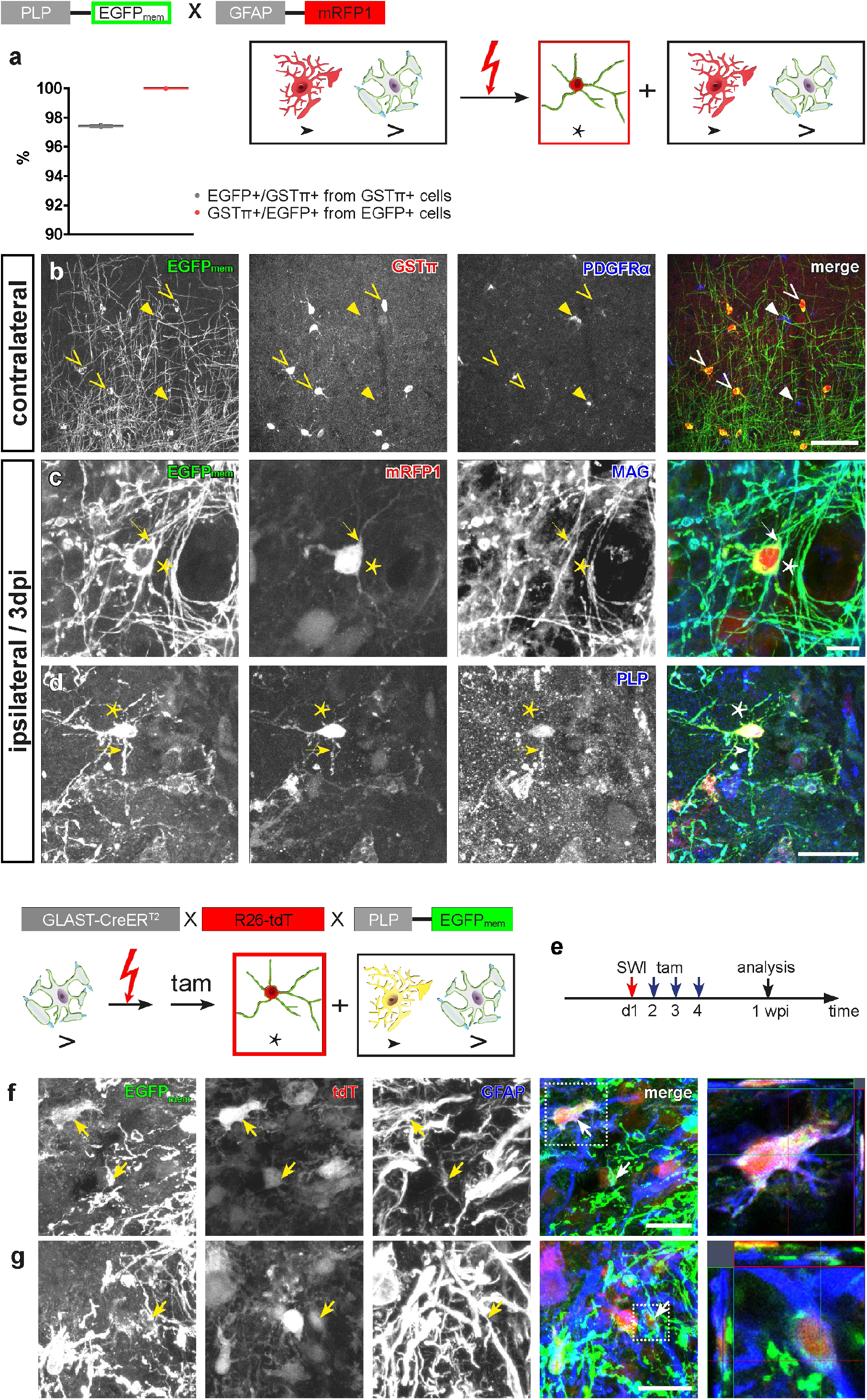
Identification of AO cells and their differentiation into astrocytes with PLP-EGFP_mem_/GFAP-mRFP1 mice. **a**, Quantification of EGFP transgene efficiency and proportion of EGFP-expressing cells. **b**, EGFP was exclusively expressed by mature oligodendrocytes (GSTπ+), but not by OPCs (PDGFRα+) in the intact cortex. **c, d**, AO cells expressed mature oligodendrocyte markers MAG (**c**) and PLP (**d**) 3 dpi. **e**, Crossbreeding of PLP-EGFP_mem_with the astrocyte-specific Cre-inducible mouse line GLAST-CreER^T2^. Tamoxifen was administrated at the second day after SWI to label oligodendrocytes with Glast locus activity. **f, g**, At 1 wpi, EGFP_mem_+/tdT+/GFAP+ cells appeared around the lesion site, indicating that oligodendrocytes do not only activate the GFAP promoter, but also other astrocytic promoters as detected for the GLAST gene. Open triangles: oligodendrocytes, triangles: OPCs, asterisks: AO cells. Scale bars in **b** = 50 µm, **c** = 10 µm, **d** = 25 µm, **f, g** = 20 µm.

**Supplementary Figure 9.**
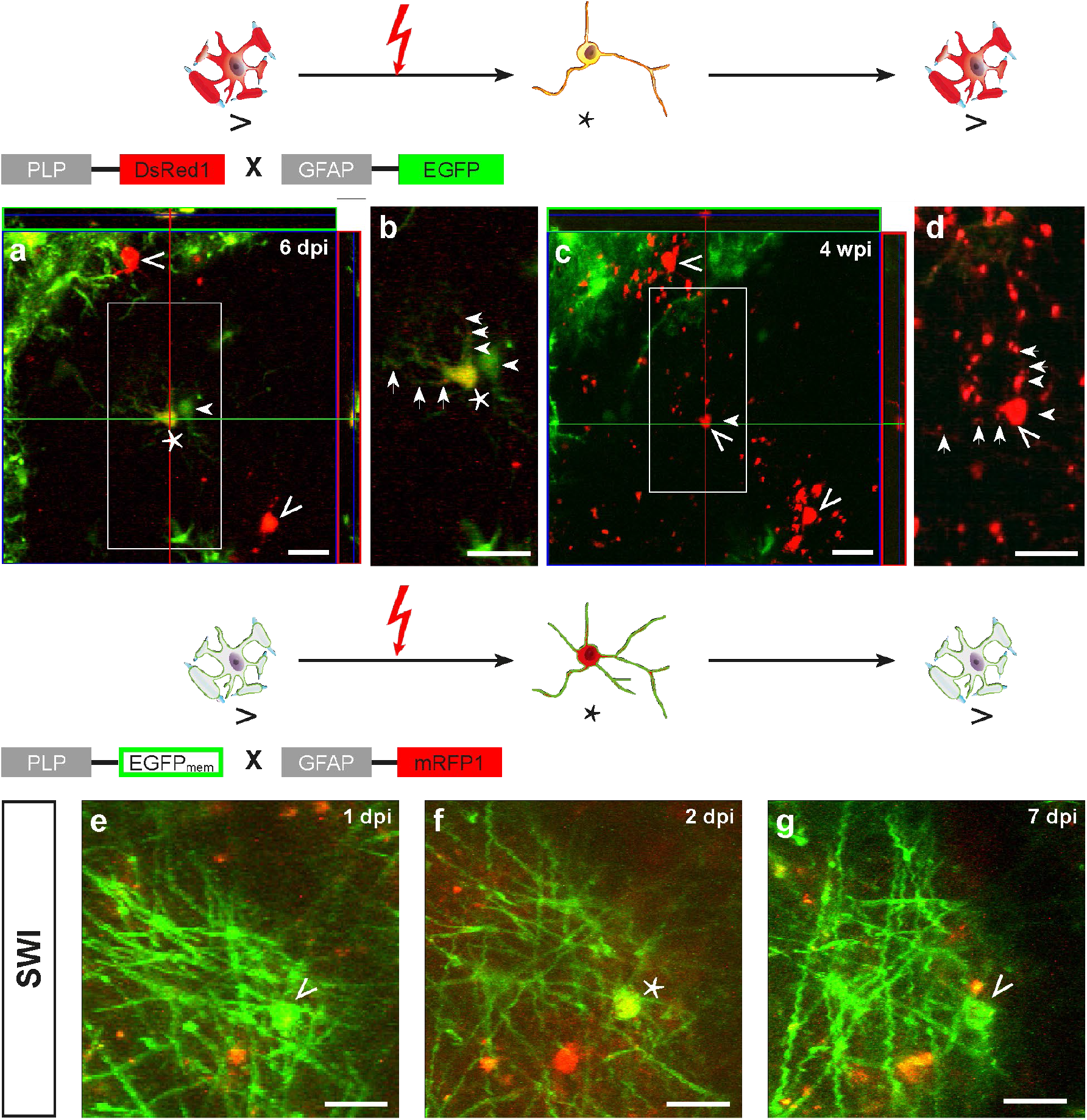
*In vivo* 2PLSM visualizes the differentiation of AO cells into oligodendrocytes. **a-d**, Three weeks of AO cell tracing in PLP-DsRed1/GFAP-EGFP mouse revealed the conversion/differentiation of an AO cell (asterisk) into an oligodendrocyte (open triangles in **a, c**). **b, d**, Magnified views of the AO cell (asterisk) that down-regulated EGFP expression in somata and processes (arrows) and differentiated into an oligodendrocyte (open triangle) four weeks after SWI. Note the disappearance of an EGFP-expressing astrocyte (arrowhead) between 6 dpi (**a, b**) and 28 dpi (**c, d**). **e-g**, An oligodendrocyte started to express mRFP1 to become an AO cell at 2 dpi (asterisk in **e**) and lost mRFP1 expression already at 7 dpi (open triangle, **g**). Scale bars = 20µm.

**Supplementary Figure 10.**
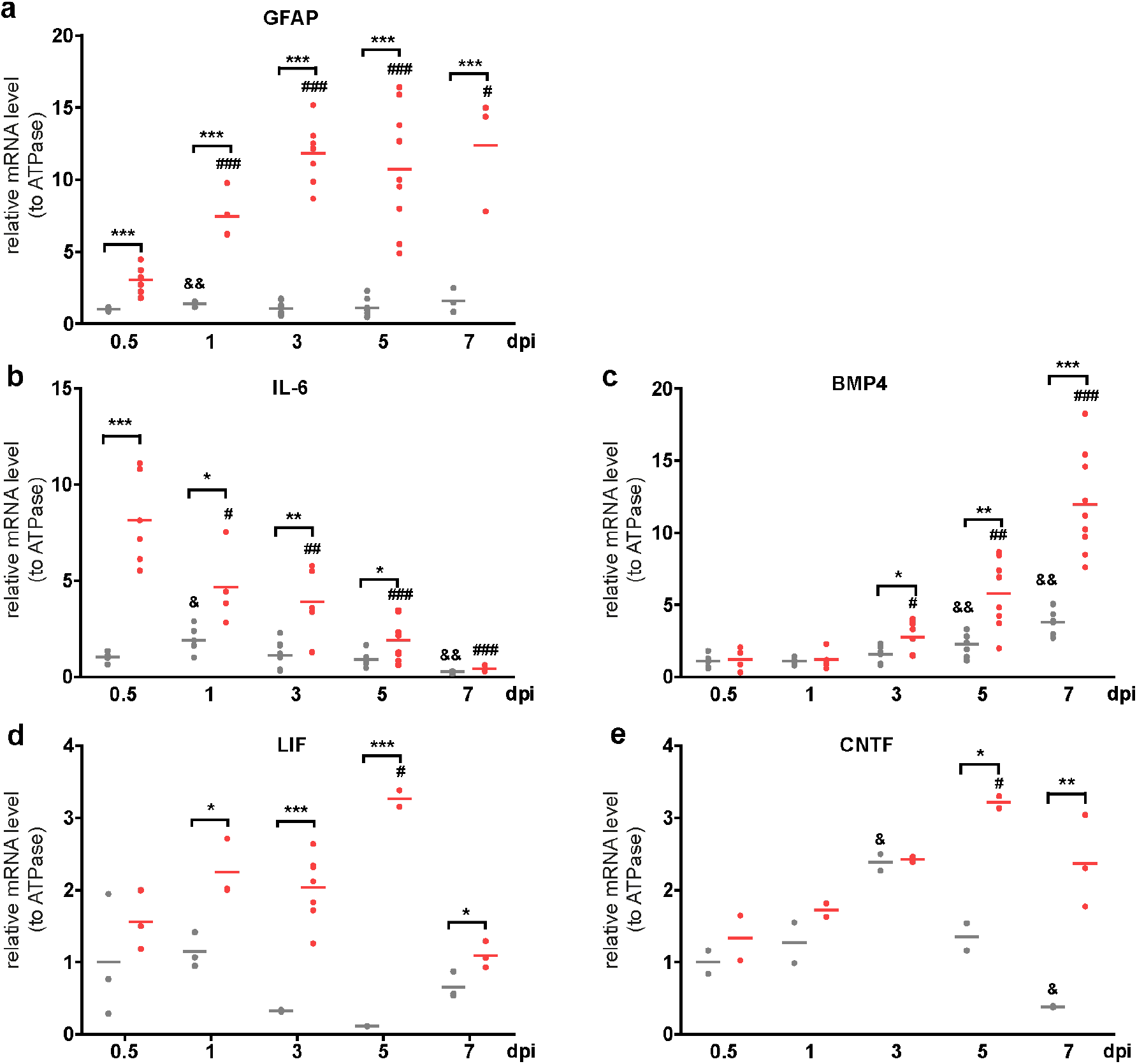
Quantification of relative mRNA levels. GFAP (**a**), IL-6 (**b**), BMP4 (**c**), LIF (**d**) and CNTF (**e**) compared to ATPase of ipsi-and contralateral cortex tissue at different time points after SWI. *: p < 0.05, **: p < 0.01 and ***: p < 0.001, compared with the corresponding contralateral side; #: p < 0.05, ##: p < 0.01 and ###: p < 0.001, compared with ipsilateral side of 0.5 dpi; &: p < 0.05 and &&: p < 0.01, compared with contralateral side of 0.5 dpi, one-way ANOVA, for detailed p-value information see **Supplementary table 4 online**.

## Supplementary Movies

**Supplementary Movie 1. Repeated, long-term *in vivo* 2P-LSM recordings over 8 days post injury**. Oligodendrocytes (DsRed1^+^) convert into AO cells (DsRed1^+^EGFP^+^) and further differentiate into astrocytes (EGFP^+^) in PLP-DsRed1/GFAP-EGFP mice. Arrows indicate two individual oligodendrocytes converting during the imaging period.

**Supplementary Movie 2. *In vivo* 2P-LSM recordings over 50 days post injury**. An oligodendrocyte converts into an AO cell and further differentiates into an astrocyte in PLP-EGFP_mem_/GFAP-mRFP1. The arrow indicates the converting oligodendrocyte.

### Abbreviations

BMP4: bone morphogenetic protein 4
BrdU: 5-bromo-2’-deoxyuridine
CNTF: ciliary neurotrophic factor
dbi: days before injury
dpi: days post injury
GLAST: glutamate/aspartate transporter
IL-6: interleukin 6
LIF: leukemia inhibitory factor
MCAO: middle cerebral artery occlusion
AO cell: astro-and oligodendroglial -precursor cell
OPC: oligodendrocyte precursor cell
R26-tdT: Rosa26Actin^fl^STOP^fl^tdTomato
SWI: stab wound injury
tam: tamoxifen
tdT: tdTomato
wbi: weeks before injury
wpi: weeks post injury
2P-LSM: two-photon laser-scanning microscopy

