## Supplementary table 1 for "After traumatic brain injury oligodendrocytes regain a plastic phenotype and can become astrocytes"

| **Depth from pia** [mm] | **Moue line** | | | | | | | |
| --- | --- | --- | --- | --- | --- | --- | --- | --- |
|  | **split-Cre x R26-tdT** | | | | | | **NG2-CT2 x R26-tdT** | |
|  | 1 wpi | | 2 wpi | | 4 wpi | |  | |
| 0 - 0.3 | GFAP^+^:  GSTπ^+^:  PDGFRα^+^:  Other: | 76.6 ± 7.5  5.7 ± 2.7  12.1 ± 4.3  5.7 ± 3.9 | GFAP^+^:  GSTπ^+^:  PDGFRα^+^:  Other: | 70.7 ± 1.9  5.7 ± 2.7  4.6 ± 1.9  19.7 ± 4.1 | GFAP^+^:  GSTπ^+^:  PDGFRα^+^:  Other: | 58.8 ± 8.7  3.6 ± 2.5  15.8 ± 6.1  26.9 ± 9.6 | GFAP^+^:  GSTπ^+^: | 78.2 ± 5.9  11.9 ± 2.8 |
| 0.3 - 0.6 | GFAP^+^:  PDGFRα^+^:  GSTπ^+^:  Other: | 55.9 ± 6.6  14.8 ± 4.9  24.1 ± 9.8  5.3 ± 3.8 | GFAP^+^:  GSTπ^+^:  PDGFRα^+^:  Other: | 58.6 ± 9.9  12.5 ± 6.7  30.3 ± 8.7*  (p=0.016, one-way ANOVA)  4.9 ± 4.3 | GFAP^+^:  PDGFRα^+^:  GSTπ^+^:  Other: | 62.9 ± 8.1  5.2 ± 3.7  17.8 ± 7.8  17.2 ± 6.5 | GFAP^+^:  GSTπ^+^: | 53.7 ± 9.4  36.9 ± 10.6 |
| 0.6 - 0.9 | GFAP^+^:  PDGFRα^+^:  GSTπ^+^:  Other: | 52.0 ± 9.4  16.4 ± 9.4  35.0 ± 7.7 ***  (p=0.0037, one-way ANOVA)  0 | GFAP^+^:  PDGFRα^+^:  GSTπ^+^:  Other: | 44.0 ± 4.2  14.6 ± 7.9  41.0 ± 5.3 ***  (p=3.45E-05,  one-way ANOVA  5.4 ± 4.6 | GFAP^+^:  PDGFRα^+^:  GSTπ^+^:  Other: | 53.9 ± 8.2  0  22.2 ± 4.8  23.9 ± 9.4 | PDGFRα^+^:  GSTπ^+^: | 44.2 ± 5.1*  (p=0.0116, one-way ANOVA)  48.1 ± 5.0 **  (p=0.003, one-way ANOVA) |

**Supplementary Table 1. Region-dependent distribution of glial cells after SWI.** Quantification of AO cell-derived and OPCs-derived glial cells in three cortical regions of split-Cre and NG2-CreER^T2^ mice after 1, 2 and 4 weeks post injury (wpi). Astrocytes: GFAP^+^; oligodendrocytes: GSTπ^+^; OPCs: PDGFRα^+^. Unidentified cells are indicated as “other”. *: p < 0.05; **: p < 0.01; ***: p < 0.001, compared to cortical region underneath the pia (0-0.3 mm) at the same time point of analysis (i.e. within a vertical column).
