## Supplementary table 2 for "After traumatic brain injury oligodendrocytes regain a plastic phenotype and can become astrocytes"

| **Abbrevation** | **Mouse line** | **Additional reporter line** | **Purpose** | **Cell type** | | **Publication** |
| --- | --- | --- | --- | --- | --- | --- |
|  |  |  |  | **Intact cortex** | **lesioned cortex** |  |
| NG2-CreER^T2^ | TgH(NG2-CreERT2)_NGCE_ | TgH(Rosa26Actin^fl^STOP^fl^  tdTomato)_R26-tdT_ | Labeling over 90 % of OPCs after one week of tamoxifen induction | tdT^+^ OPCs;  tdT^+^ oligodendrocytes | tdT^+^ OPCs;  tdT^+^ oligodendrocytes;  **tdT^+^ astrocytes** | ^1, 2^ |
| NG2-CreER^T2^  /  GFAP-EGFP | TgH(NG2-CreERT2)_NGCE_  x  TgN(hGFAP-EGFP)_GFEC_ | TgH(Rosa26Actin^fl^STOP^fl^  tdTomato)_R26-tdT_ | Observation of OLCs activation after acute cortical injuries;  EGFP expression by 38.5 % of astrocytes in GFEC mouse line | tdT^+^ OPCs;  tdT^+^ oligodendrocytes;  EGFP^+^ astrocytes | tdT^+^ OPCs;  tdT^+^ oligodendrocytes;  EGFP^+^ astrocytes;  **tdT^+^EGFP^+^ OPCs;**  **tdT^+^EGFP^+^ AO cells** | ^1-3^ |
| split-Cre | TgN(hGFAP-NCre)_GCNT_  x  TgN(mPLP-CCre)_PCCK_ | TgH(Rosa26^fl^STOP^fl^EYFP)_R26-EYFP_ or  TgH(Rosa26Actin^fl^STOP^fl^ tdTomato)_R26-tdT_ | 1) Exclusively labeling the AO cells without activated OPCs    2-3) Investigation of stability of   split-Cre construction | n.d. | **reporter^+^ AO cells;**  **reporter^+^ AO cell-derived cells** | ^2, 4, 5^ |
| PLP-DsRed1/  GFAP-EGFP | TgN(hGFAP-EGFP)_GFEA_  x  TgN(mPLP-DsRed1)_PRDB_ | - | Observation and characterization of the AO cells; EGFP expression by 42.7 % of astrocytes in GFEA mouse line | EGFP^+^ astrocytes;  9 % of EGFP^+^ OPCs;  DsRed1^+^ oligodendrocytes | EGFP^+^ astrocytes;  DsRed1^+^ oligodendrocytes;  **EGFP^+^DsRed1^+^ AO cells** | ^6, 7^ |
| PLP-EGFP_mem_  /  GFAP-mRFP1 | TgN(mPLP-EGFP_mem_)  x  TgN(hGFAP-mRFP1) |  | Observation and characterization of the AO cells; 100 % of EGFPmem expression by oligodendrocytes | EGFP^+^ oligodendrocytes;  mRFP1^+^ astrocytes | EGFP^+^ oligodendrocytes;  mRFP1^+^ astrocytes;  **EGFP^+^mRFP1^+^ AO cells** | ^8, 9^ |
| Glast-CreER^T2^/  PLP-EGFP_mem_ | TgN(PLP-EGFPmem)_PLPG_  x  TgH(Glast-CreERT2)_GLAC_ | TgH(Rosa26Actin^fl^STOP^fl^  tdTomato)_R26-tdT_ | Observe AO cells with another astrocyte specific transgenic mouse | EGFP^+^ oligodendrocytes;  tdT^+^ astrocytes | EGFP^+^ oligodendrocytes;  tdT^+^ astrocytes;  **EGFP^+^tdT^+^ AO cells** | ^6^ |
| NG2-EYFP | TgH(NG2-EYFP)_NGYF_ | - | Exclusively labeling of OPCs | EYFP^+^ OPCs | EYFP^+^ OPCs | ^10^ |
| GFAP-CreER^T2^ | TgN(hGFAP-CreERT2)_GCTF_ | TgH(Rosa26^fl^STOP^fl^EYFP) _R26-EYFP_ | Labeling cells derived from astrocyte lineage | EYFP^+^ astrocytes | EYFP^+^ astrocytes | ^5, 11^ |
| PLP-CreER^T2^ | TgN(mPLP-CreERT2)_PCET_ | TgH(Rosa26^fl^STOP^fl^EYFP) _R26-EYFP_ | Labeling cells derived from oligodendrocyte lineage | EYFP^+^ oligodendrocytes;  EYFP^+^ OPCs | EYFP^+^ oligodendrocytes;  EYFP^+^ OPCs;  EYFP^+^ astrocytes | ^12, 13^ |

**Supplementary Table 2. General description of mouse lines.** The transgenic mouse lines of this study have been generated previously by homologous and non-homologous gene recombination. Conditional transgenic mice carrying split-Cre or CreER^T2^ DNA recombinase were bred with additional reporter mice to visualize recombined cells. In the intact and injured cortex, different populations of cell types were observed as listed in the table. n.d.: not detectable.

**References**
