## Supplementary table 3 for "After traumatic brain injury oligodendrocytes regain a plastic phenotype and can become astrocytes"

| **Mouse line** | **Cell type** | **Area** [mm^2^] | **Depth from pia** [mm] | **Width from lesion** (0,[mm]) |
| --- | --- | --- | --- | --- |
| PLP-DsRed1/GFAP-EGFP | AO cell | 0.6 | 0 - 1 | -0.3 - 0.3 |
| split-Cre | AO cell-derived recombined cells | 0.6 | 0 - 1 | -0.3 - 0.3 |
| NG2-CreER^T2^ x R26-tdT | tdT+ recombined cells | 0.6 | 0 - 1 | -0.3 - 0.3 |
| split-Cre | AO cell-derived recombined cells (Region dependent distribution) | 0.18  (neuronal layer I-III) | 0 - 0.3 | -0.3 - 0.3 |
|  |  | 0.18  (neuronal layer IV-V) | 0.3 - 0.6 | -0.3 - 0.3 |
|  |  | 0.18  (neuronal layer VI) | 0.6 - 0.9 | -0.3 - 0.3 |

**Supplementary Table 3 Region-selection for cell counting**. Cells within 300 µm left and right from the lesion site were counted down to 1 mm cortex depth. In the region-dependent differentiation study of split-Cre mice, the cortex was subdivided into three subgroups; each of them was 300 µm in depth.
