## Supplementary table 4 for "After traumatic brain injury oligodendrocytes regain a plastic phenotype and can become astrocytes"

|  |  | il 0.5 | cl 0.5 | cl 1 | cl 3 | cl 5 | cl 7 |
| --- | --- | --- | --- | --- | --- | --- | --- |
| **GFAP**  **mRNA**  **(Supplementary Fig. 13a)** | **cl 0.5** | p=0.0005, *** |  | p=0.0032, && | p=0.816, n.s. | p=0.684, n.s. | p=0.103, n.s |
|  | **il 0.5** |  | p=0.0005, *** | n.c. | n.c. | n.c. | n.c. |
|  | **il 1** | p=0.0007, ### | n.c. | p=0.0004, *** | n.c. | n.c. | n.c. |
|  | **il 3** | p=3.95E-07, ### | n.c. | n.c. | p=8.61E-11, *** | n.c. | n.c. |
|  | **il 5** | p=0.0008, ### | n.c. | n.c. | n.c. | p=1.27E-05, *** | n.c. |
|  | **il 7** | p=0.0007, ### | n.c. | n.c. | n.c. | n.c. | p=0.0102, * |
| **IL-6**  **mRNA**  **(Supplementary Fig. 13b)** | **cl 0.5** | p=0.0004, *** |  | p=0.0375, & | p=0.829, n.s. | p=0.616, n.s. | p=0.007, && |
|  | **il 0.5** |  | p=0.0004, *** | n.c. | n.c. | n.c. | n.c. |
|  | **il 1** | p=0.0424, # | n.c. | p=0.0132, * | n.c. | n.c. | n.c. |
|  | **il 3** | p=0.0096, ## | n.c. | n.c. | p=0.00229, ** | n.c. | n.c. |
|  | **il 5** | p=2.47E-05, ### | n.c. | n.c. | n.c. | p=0.043, * | n.c. |
|  | **il 7** | p=0.00094, ### | n.c. | n.c. | n.c. | n.c. | p=0.159, n.s. |
| **BMP4**  **mRNA**  **(Supplementary Fig. 13c)** | **cl 0.5** | p=0.737, n.s. |  | p=0.994, n.s. | p=0.125, n.s. | p=0.006, && | p=5.01E-06, &&& |
|  | **il 0.5** |  | p=0.737, n.s. | n.c. | n.c. | n.c. | n.c. |
|  | **il 1** | p=0.969, n.s. | n.c. | p=0.68, n.s. | n.c. | n.c. | n.c. |
|  | **il 3** | p=0.037, # | n.c. | n.c. | p=0.0188, * | n.c. | n.c. |
|  | **il 5** | p=0.0051, ## | n.c. | n.c. | n.c. | p=0.0016, ** | n.c. |
|  | **il 7** | p=9.92E-05, ### | n.c. | n.c. | n.c. | n.c. | p=3.83E-06, *** |
| **LIF**  **mRNA**  **(Supplementary Fig. 13d)** | **cl 0.5** | p=0.364, n.s. |  | p=0.793, n.s. | p=0.244, n.s. | p=0.154, n.s. | p=0.534, n.s. |
|  | **il 0.5** |  | p=0.364, n.s. | n.c. | n.c. | n.c. | n.c. |
|  | **il 1** | p=0.108, n.s. | n.c. | p=0.015, * | n.c. | n.c. | n.c. |
|  | **il 3** | p=0.166, n.s. | n.c. | n.c. | p=0.00003, *** | n.c. | n.c. |
|  | **il 5** | p=0.0125, # | n.c. | n.c. | n.c. | p=4.35E-05, *** | n.c. |
|  | **il 7** | p=0.147, n.s. | n.c. | n.c. | n.c. | n.c. | p=0.045, * |
| **CNTF**  **mRNA**  **(Supplementary Fig. 13e)** | **cl 0.5** | p=0.44, n.s. |  | p=0.495, n.s. | p=0.02, & | p=0.294, n.s. | p=0.0146, & |
|  | **il 0.5** |  | p=0.44, n.s. | n.c. | n.c. | n.c. | n.c. |
|  | **il 1** | p=0.357, n.s. | n.c. | p=0.267, n.s. | n.c. | n.c. | n.c. |
|  | **il 3** | p=0.074, n.s. | n.c. | n.c. | p=0.765, n.s. | n.c. | n.c. |
|  | **il 5** | p=0.0281, # | n.c. | n.c. | n.c. | p=0.012, * | n.c. |
|  | **il 7** | p=0.145, n.s. | n.c. | n.c. | n.c. | n.c. | p=0.0057, ** |

**Supplementary Table 4 Exact p-values of comparative analysis of RT-PCR results.** Quantification of RT-PCR results of different time points after injury (0.5, 1, 3, 5 and 7 d) and of contralateral (cl) and ipsilateral (il) side as shown in Supplementary Figure 12 *: p < 0.05, **: p < 0.01 and ***: p < 0.001, compared with the corresponding contralateral side; #: p < 0.05, ##: p < 0.01 and ###: p < 0.001, compared with ipsilateral side of 0.5 dpi; &: p < 0.05 and &&: p < 0.01, compared with contralateral side of 0.5 dpi, one-way ANOVA. n.c.: not compared
